# Stress generation, relaxation and size control in confined tumor growth

**DOI:** 10.1101/761817

**Authors:** Huaming Yan, Daniel Ramirez-Guerrero, John Lowengrub, Min Wu

**Affiliations:** Department of Mathematics, University of California, Irvine, Irvine, California, USA; Center for Multiscale Cell Fate Studies, University of California, Irvine, Irvine, California, USA; Department Biomedical Engineering, University of California, Irvine, Irvine, California, USA; Center for Complex Biological Systems, University of California, Irvine, Irvine, California, USA; Department of Mathematical Sciences, Worcester Polytechnic Institute, Worcester, Massachusetts, USA

## Abstract

Experiments on tumor spheroids have shown that compressive stress from their environment can reversibly decrease tumor expansion rates and final sizes. Stress release experiments show that nonuniform anisotropic elastic stresses can be distributed throughout. The elastic stresses are maintained by structural proteins and adhesive molecules, and can be actively relaxed by a variety of biophysical processes. In this paper, we present a new continuum model to investigate how the growth-induced elastic stresses and active stress relaxation, in conjunction with cell size control feedback machinery, regulate the cell density and stress distributions within growing tumors as well as the tumor sizes in the presence of external physical confinement and gradients of growth-promoting chemical fields. We introduce an adaptive reference map that relates the current position with the reference position but adapts to the current position in the Eulerian frame (lab coordinates) via relaxation. This type of stress relaxation is similar to but simpler than the classical Maxwell model of viscoelasticity in its formulation. By fitting the model to experimental data from two independent studies of tumor spheroid growth and their cell density distributions, treating the tumors as incompressible, neo-Hookean elastic materials, we find that the rates of stress relaxation of tumor tissues can be comparable to volumetric growth rates. Our study provides insight on how the biophysical properties of the tumor and host microenvironment, mechanical feedback control and diffusion-limited differential growth act in concert to regulate spatial patterns of stress and growth. When the tumor is stiffer than the host, our model predicts tumors are more able to change their size and mechanical state autonomously, which may help to explain why increased tumor stiffness is an established hallmark of malignant tumors.

**Author summary:** The mechanical state of cells can modulate their growth and division dynamics via mechanotransduction, which affects both the cell size distribution and the tissue size as a whole. Experiments on tumor spheroids have shown that compressive stress from their environment can reversibly decrease tumor expansion rates and final sizes. Besides external confinement and compression on the tumor border, a heterogeneous stress field can be generated inside the tumor by nutrient-driven differential growth. Such growth-induced mechanical stresses can be relaxed by tissue rearrangement, which happens during cell neighbor exchanges, cell divisions, and extracellular matrix renewal. In this study, we have developed a continuum model that describes the above mechanical interactions and the dynamics of tissue rearrangement explicitly. Motivated by published experimental data, we consider mechanotransduction where the local compressive stress slows down cell growth and cell size reduction limits cell division. We have analyzed how external mechanical stimuli and internal processes influence the outcome of cell-and-tissue sizes and spatial patterns of cell density and mechanical stress in growing tumors.

## Introduction

The importance of mechanical forces in regulating cell behaviors during normal development and homeostasis [1,2], and in disease progression such as cancer [3–5], is now widely recognized. In addition to driving cell movement and deformation, biophysical forces act via mechanotransduction to regulate cell fates, division, and death rates [6–9]. Nevertheless, the effects of tissue properties and mechanical stresses on cancer progression and treatment outcomes are still not well understood. *In vitro* experiments on avascular tumor spheroids have shown that changes in their mechanical environment can result in different tumor expansion rates, stress distributions, final sizes [10–13], and altered cell density fields [13]. Further, it was recently observed that the material properties of tissues and external mechanical compression may play distinct roles in tumor growth [14]. In addition to mechanical compression from the exterior environment, stresses within growing tumors can be generated by nutrient-driven differential growth of cells as well as active contractility of cells [15]. The release of these stresses, by cutting, slicing, or punch protocols [14, 16], can cause finite deformations at rates faster than those of a typical cell cycle and these deformations can be used to estimate the residual elastic stress. The elastic stresses are maintained by structural proteins and adhesive molecules [17], and can be actively relaxed due to processes such as turnover/reassembly of structural and adhesion molecules, cell re-arrangements, and oriented cell divisions [1] at timescales spanning from minutes to hours [17, 18].

In this paper, we present a new continuum model in the laboratory (Eulerian) frame. This approach facilitates the nonlinear coupling between finite-elastic stresses, which are traditionally described using material coordinates (Lagrangian frame), and reaction-diffusion equations that model tissue growth and the dynamics of cell substrates (e.g, oxygen and nutrients), which are traditionally described in the Eulerian frame. We fit the model to experimental data from two independent studies of tumor spheroid growth and perform parametric studies around the fitted parameter regime. We investigate how a tumor spheroid modulates its size as a whole, as well as cell density and stress distribution via internal stress relaxation in the context of external physical confinement and limited nutrient diffusion in conjunction with feedback machineries on cell growth [19–21] and division [22,23].

## Methods

### Tissue elasticity in an Eulerian frame

We formulate the elastic stresses using Eulerian coordinates. Instead of mapping the reference coordinates of a material point to the current coordinates as done in a Lagrangian frame, we introduce the reference map *y*(*r,t*), which maps the current radial coordinates of a material point to its reference radial position in the tissue. Treating the tumor spheroid as an incompressible, neo-Hookean elastic material the nondimensional elastic stresses can be written as

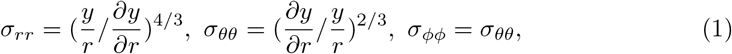

where we have nondimensionalized the stress using the shear modulus of the tumor tissue. This approach can also be used for more general constitutive laws [24]. The total stress (Cauchy stress tensor) is *σ^tot^* = ***σ*** – *p***I** where *p* is the pressure. The tumor tissue is approximately incompressible in its elastic response since most of its volume fraction is water. However, volume loss can occur against compression due to water efflux [13], which we will introduce shortly. Further, time and space are nondimensionalized using the characteristic cell-cycle time scale of *τ* = 1 day and a length scale of *l* = 1*μm*. See Text S1 for the full, nonsymmetric, dimensional model and nondimensionalization.

### Adaptive reference map and stress relaxation

We describe the relaxation of the elastic stresses in Eq. (1) by introducing the relaxation rate *β* that reflects how rapidly the reference coordinates adapt to the current coordinates:

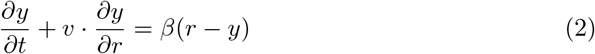

where *v* is the cell velocity and *y*(*r*, 0) = *r*, assuming the initial state is unstressed. When *β* = 0, *y* satisfies *∂_t_y*(*r,t*) + *v*(*r,t*)*∂_r_y*(*r,t*) = 0, which means the reference coordinate of a material point is unchanged along its trajectory [25,26]. In the results, we will show a positive value of *β* (adaptation) is necessary for the mechanical equilibrium when the tumor reaches its equilibrium size. Notice the dynamics of the displacement field *u* = *r* – *y* becomes 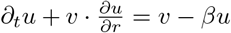, which decays with a rate of *β*. Combining Eqs. (1) and (2) yields a Maxwell-like stress relaxation. See Text S1 for more details. This is similar in spirit to the approach used in [27] to model viscoelasticity in the context of a model of the cell cytoskeleton.

### Force balance inside the tumor

We model the tissue as an overdamped, incompressible, nonlinear elastic material so that at any point in the tissue the velocity is proportional to the force. Using radial symmetry, the nondimensional system is:

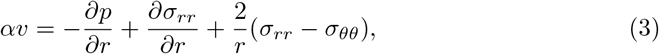

where *α* is the nondimensional friction (drag) coefficient and *v* is the radial velocity. We denote the total radial and circumferential stresses by 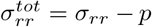 and 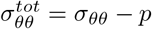, respectively.

### External compression

At the spheroid boundary *r* = *R*, we have the force balance relation

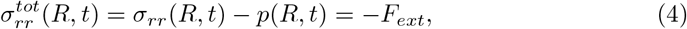

where –*F_ext_* represents the result of external physical confinement from a surrounding solid material, such as externally applied hydrostatic pressure, or surface tension. Assuming that the spherical tissue grows within an incompressible neo-Hookean material, similar to the experimental set-up in [10], the external compression at the tumor boundary can be written as a function of the initial and current radius, *R*_0_ and *R*(*t*), respectively (see Section 1.3 in Text S1):

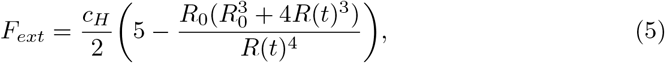

where *c_H_* is the shear modulus of the surrounding material relative to that of the tumor tissue. Alternatively, if a hydrostatic pressure 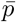 is applied, as in [12,13], then 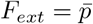.

### Cell density

We assume the nondimensional local cell proliferation rate is given by 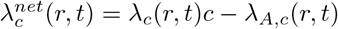, where *λ_c_*(*r, t*) is the rate of cell divisions, *c* is the concentration of a growth promoting biochemical factor that represents the net effect of diffusible substances (e.g., oxygen, glucose) on cell division, and *λ_A,c_*(*r, t*) is the rate of apoptosis. Henceforth, the growth promoting factor *c* is referred to as nutrient.

The cell number density *ρ_c_* (cell numbers per unit volume) should follow

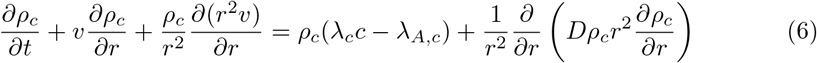

where 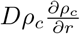 models the cell number flux due to local cell neighbor exchanges. When 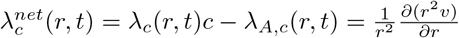, we have *ρ_c_* ≡ *const*.. In general, it is not the case, and the rates of cell division and apoptosis and the rates of cell volumetric growth and loss (introduced below) are not necessarily matched, respectively.

### Volumetric growth

We assume that the rate of local volume change is given by *λ^net^*(*r, t*) = λ(*r, t*)*c* – λ_*A*_(*r, t*), where λ(*r, t*) is the rate of volume increase (e.g., cell volume growth) and λ_*A*_(*r, t*) = λ_*A,c*_ + λ_*E*_ is the rate of volume loss from cell apoptosis and the rate of water efflux λ_*E*_. Considering that the majority of its volume fraction is water, the tumor spheroid is incompressible such that 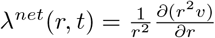. Thus the radial velocity *v*(*r,t*) is:

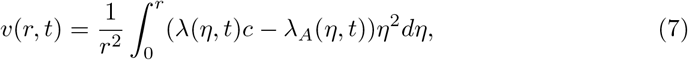

and the tumor radius evolves via *dR*(*t*)/*dt* = *v*(*R, t*). As the tumor size changes, *c* can be modeled as

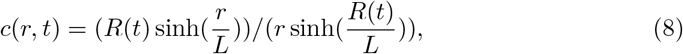

assuming the substrate diffuses in from the tissue boundary, with *c*(*R*(*t*), *t*) = 1, and is uptaken at a constant rate by tumor cells. Here, *L* is the nondimensional diffusion length. Note that *c*(*R*(*t*), *t*) could be a function of time, but we assume here that there is sufficient supply from the microenvironment to maintain *c*(*R*(*t*),*t*) constant. See Section 1.3 in Text S1 for details.

### Water efflux and mechanotransduction

In growing tumor spheroids, compressive stresses can inhibit cell proliferation (e.g., [12,13]) as well as inducing water efflux [13]. Thus, we assume that local compressive stress induces local cell water efflux (with rate λ_*E*_) and inhibits cell volume growth (with rate λ). We model the effect of compression on water flux and cell volume growth by

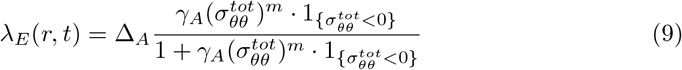

and

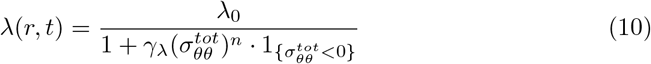

respectively, as Hill-type equations where is the base rate of cell volume growth, *γ_A_* and *γ_A_* are feedback strengths, Δ_*A*_ is the maximum rate of the water flux, *m* and *n* are positive even integers, and 1_χ_ denotes the characteristic function of the set χ. Eq. (9) takes into the account of Michaelis-Menten kinetics with a saturation effect when the local compressive stress becomes large in magnitude. The cell growth rate (Eq.(10)) decreases from the baseline rate λ_0_ in response to the local compressive stress. We also assume that circumferential stresses 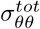 dominate the mechanical state of the cells because the circumferential stress represents stresses from two principal directions while radial stress 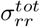 accounts for only one. In the Results section, we validate this assumption (see *Inhibition of growth through external confinement*). The water efflux process may require a detailed consideration of flux exchange with the tumor interstitial space. For example, it is possible that the interstitial pressure and its associated fluid flow can limit the water efflux process from cells. For simplicity, we did not model fluid mechanics in the interstitial space. We assume that the interstitial volume fraction inside the tumor is constant and the dynamics of the water therein is slaved to the dynamics of cell growth, apoptosis, and water efflux.

Both feedbacks on λ and λ_*E*_ can result in increased cell density *ρ_c_* via Eq. (6). When cell density is high, which means the local size of cells is small, cell division can be inhibited by multiple mechanisms [22, 23]. We model this effect by

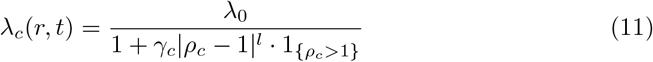

where γ_*c*_ is the feedback strength and *l* is the Hill-coefficient. We assume that the base rate of cell division is the same as the base rate of cell volume growth λ_0_. That is, without compression, the rate of cell division and the rate of cell volume growth are matched.

## Results

### A tumor spheroid growing with stress relaxation

We first simulate numerically (see Section 1.3 in Text S1 for the algorithm and parameters), the unconfined (free) growth of an initially unstressed tumor spheroid (*F_ext_* = 0). As seen in Figure 1A the tumor radius increases over time and approaches a steady-state, which is independent of the initial size or initial stress state (Figure S2 in Text S1). Because of diffusion-limited nutrient transport, the net volume rate of change *λ^net^*(*r, t*) = λ(*r, t*)*c*(*r, t*) – λ_*A*_(*r, t*) is spatially varying (Figure 1B). At early times when the tumor is small, the volume increases all throughout the tumor spheroid as nutrients are readily available. At later times, volume gain (due to cell growth) dominates at the spheroid boundary and volume loss (due to cell apoptosis and water efflux) dominates at the spheroid center (Figure 1B) where nutrient levels are low. Correspondingly, cells move outward at early times but at late times, as in previous models (e.g., [12,28,29]), cells divide at the boundary and move inward to compensate for the loss of volume at the center (Figure 1C), which may explain the presence of long-lasting apoptotic markers in the core of the tumor spheroids. The total circumferential stresses 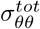 are compressive throughout the tumor spheroid at early times while at later times the stresses are compressive near the tumor edge and tensile in the center (Figure 1D). Further, the stresses equilibrate as soon as the tumor radius reaches equilibrium. When the tumor reaches its equilibrium size *R*(*t*) = *R*_∞_, one can derive *y_r_*(*R*_∞_) = *β*/ (*β* + λ(*R*_∞_) – λ_*A*_(*R*_∞_)). This shows that (i) *β* = 0 (no relaxation) leads to a singularity in the elastic strain and (ii) increasing *β* restores the proportionality between *y*(*r,t*) and *r* and thus decreases the elastic energy (Figure S3 in Text S1) and stress anisotropy (Figure S4 in Text S1), which are both 0 when *y*(*r, t*) = *r*. Similar considerations hold at the tumor center. The cell density *ρ_c_*(*r,t*) is highest in regions near the tumor boundary (Figure 1E). This is due to the synergistic effect among the rate of cell division 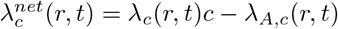, the rate of net local volume gain *λ^net^*(*r, t*) = λ(*r, t*)*c*(*r, t*) – λ_*A*_(*r, t*), and the local flux of cells. One can see from Eq. (6) that cell division tends to increase the cell density, while the local volume gain 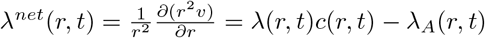 tends to decrease the cell density. When this two rates match, the cell density is uniform. However, when the compression near the tumor boundary slows down the local volume gain via mechanotransduction and water efflux, the local division rate becomes faster than that of the volume gain, which leads to an increase in cell number density. Notice the advection 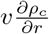 and local random neighbor exchange of cells 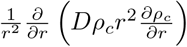 can further adjust the cell density distribution. In the central region with inward radial velocity *v*(*r, t*) < 0, local cell density increases due to the advection of the cells. At the tumor center with *v*(0, *t*) ≡ 0, the random neighbor exchange of cells 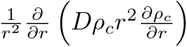 is necessary to describe local cell accumulation. This higher-order derivative term allows us the implement the no-flux boundary condition 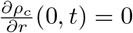 at the tumor center, which facilitates changes in cell density at the tumor center according to the cell density in its neighborhood. Interestingly, we have found the value of *D* does not visibly affect the result in the cell density distribution (see Figure S5 in Text S1). Nevertheless, the effect from the cell flux is secondary to the competition between volume gain and cell proliferation rates (in this case), due to the high proliferative activity near the free tumor boundary.

**Fig 1.**
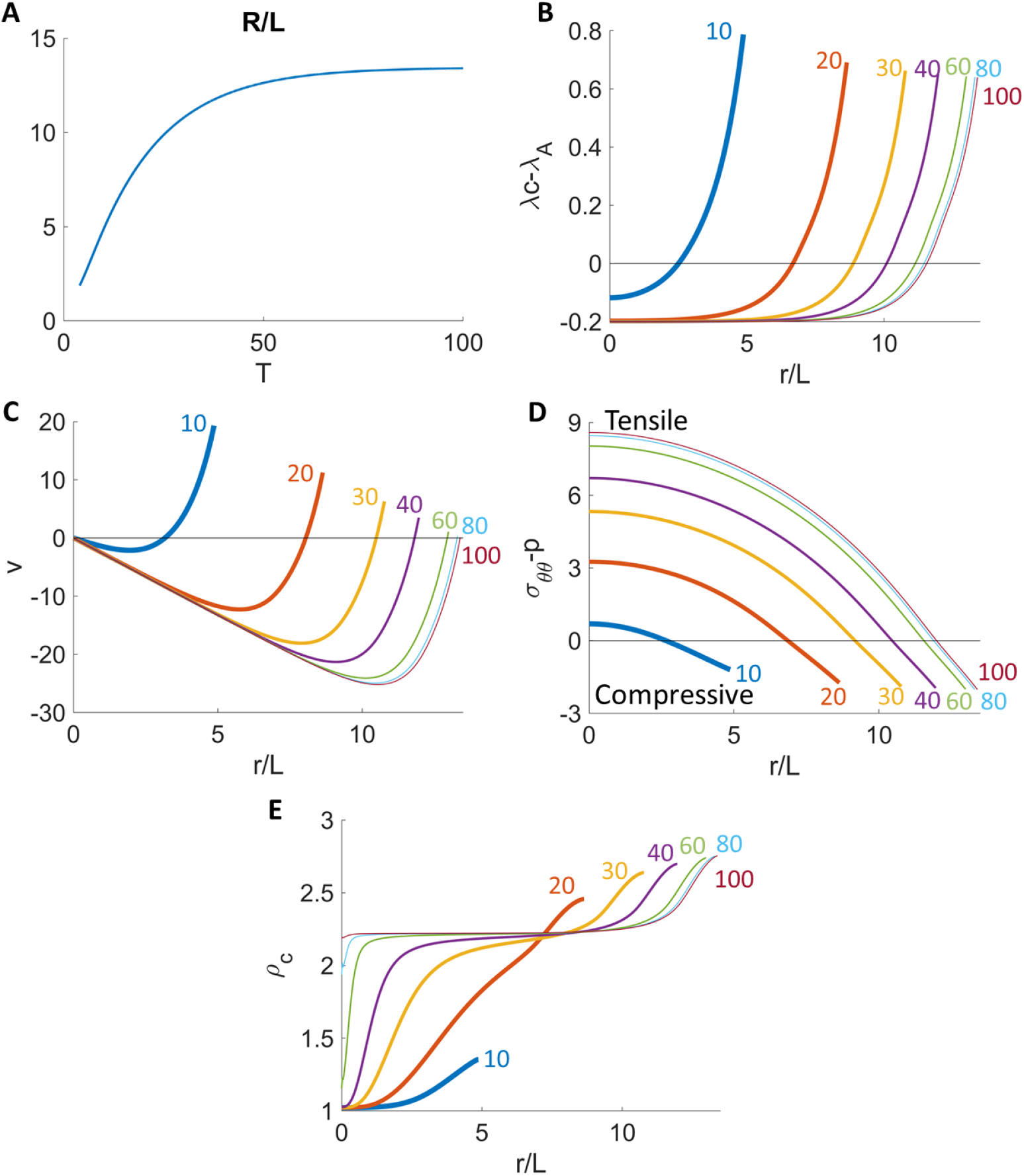
Time evolution of a freely growing tumor without external compression (A). The spatial distribution of net volumetric growth rate (B), cell velocity (C), circumferential stresses (D), and cell density (E) at different times. See Text S1 for the list of parameters.

### Inhibition of growth through external confinement

In [10], it was shown that the growth capacity of tumor spheroids (human colon adenocarcinoma, LS174T) in agarose gels decreases as the concentration of agarose is increased (Figure 2A and Figure S1 in Text S2); the stiffness of the gels is positively correlated with the agarose concentration. However, tumors suspended in gels with lower growth rates regain their free-growth capacity once the gels are removed (Figure 2E, symbols). We use our model to fit the experimental data from tumors grown in free suspension (0% gel) and 0.7% and 1.0% agarose gels using the same set of the tumor-associated parameters (which characterize the base rates and chemomechanical responses of LS174T) but different shear moduli of the gel (e.g., *c_H_* =0 for 0% and *c_H_* > 0 for the 0.7% and 1.0% gels). The experimental data consists of both tumor radius (shown as symbols in Figure 2A), and average cell densities of compressed tumors as a ratio to that of the free tumor (shown as vertical line segments denoting the mean and standard deviations of experimental measurements in Figure 2D). Since [10] suggests that cells does not adjust their rate of proliferation in response to the spatial confinement, we assume compressive stresses increase local water efflux (γ_*A*_, Δ_*A*_ > 0 and *γ*_λ_ = 0). Then, using the same tumor-associated parameters, we fit the other gel concentrations (Figure S1 in Text S2) by changing only *c_H_*. See Text S2 for details on the fitting process.

**Fig 2.**
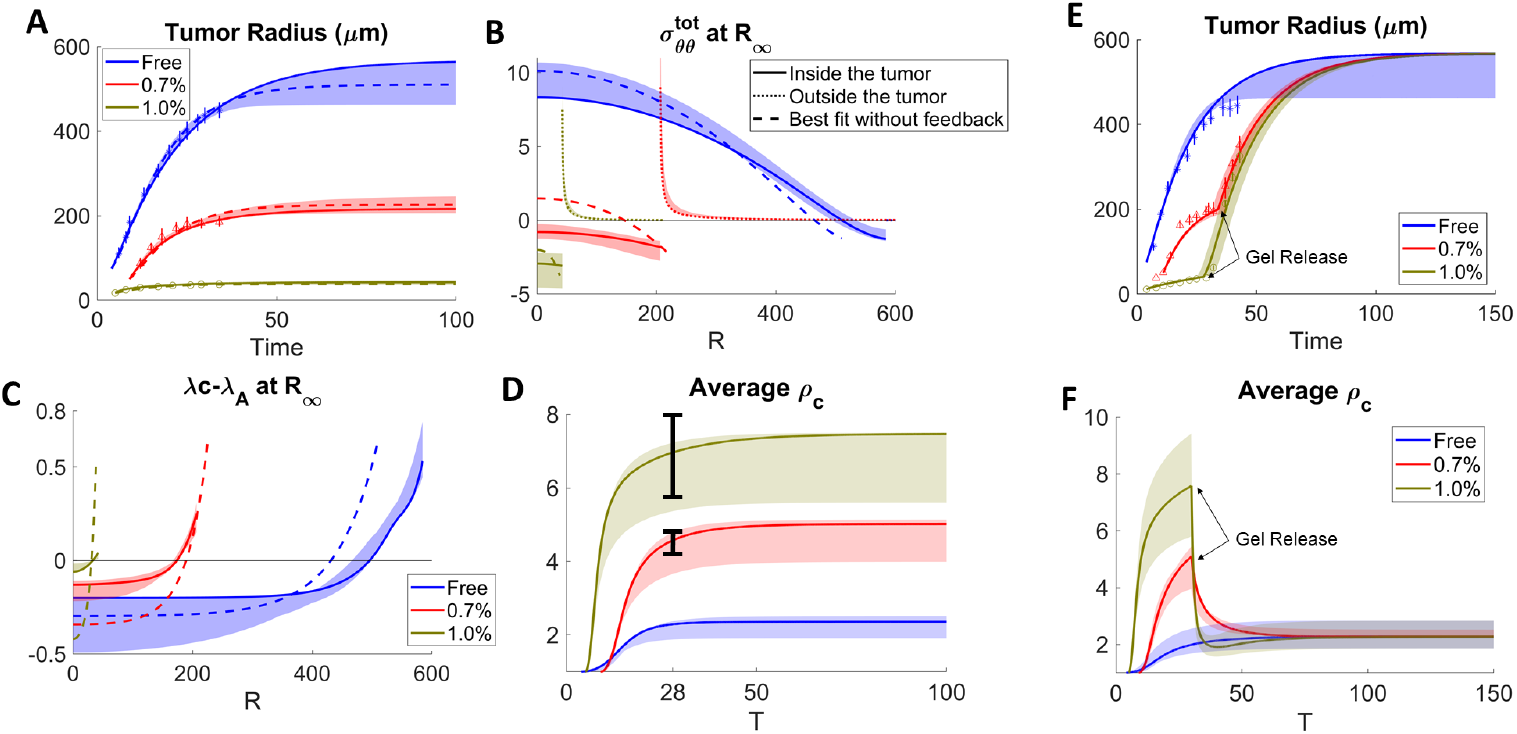
Fitting the model to data from [10] of tumor spheroids grown in free suspension (blue) and 0.7% (red) and 1.0% (brown) agarose gels with (solid) and without (dashed) feedback from the elastic stress (A) and when stresses are released by removing the gels in a narrow interval around the times reported in [10] (E). Bands show the results within 10% of the best fit with feedback in all panels. (B) and (C): The distributions of stresses and net volume growth rates from the model. (D) and (F): Average cell density of tumors in (A) and (E) respectively. The bars in (D) shows the experimental average density for 0.7% and 1.0% tumors, relative to the free tumor. See Text S2 for details.

We find that the nondimensional relaxation rate *β* ~ 1, which suggests that a fully nonlinear elastic model is needed to describe tumor biomechanics, rather than a fluid model or a linear elastic model. Such reduced models arise as limits of our model where *β* >> |λ^*net*^|. See Text S2 for limiting cases in different parameter regimes. The results are presented in Figure 2 (and Figure S1 in Text S2). There is good quantitative agreement between the model (curves) and experiments (symbols) for the dynamics of the tumor spheroid radii (Figure 2A) where the bands show the results using parameters for which the results are within 10% of the best fit, which corresponds to *β* = 0.4 and 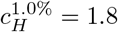 and 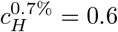 (see Text S2 for all the fitted parameters). Fits to other gel concentrations are shown in Figure S1 in Text S2. For the case of the 1. 0% gel at equilibrium, we predict the circumferential stress 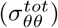 in the tumor at equilibrium is compressive and quasi-uniform (Figure 2C). We find that the average elastic energy in the tumor spheroid 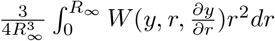 decreases even though *c_H_* increases (Figure S2 in Text S2), which shows that stress and stiffness are not always positively correlated. This result is consistent with the findings in [14]. Our results suggest that as *c_H_* increases, the stresses in the tumor become mainly hydrostatic (see Figure S2 in Text S5). When *c_H_* >> 1, it can be seen analytically that the leading-order stress distribution is uniform and is given by the compression at the boundary. As the agarose concentration of the gel is decreased, *c_H_* decreases and the stresses are less compressive and less uniform (Figure 2B, solid curves). In the freely growing case, the stress becomes tensile in the tumor interior and is compressive only at the spheroid boundary similar to that observed in Figure 1D. In contrast, the stresses in the gel (dotted curves) are tensile, as the growth of the tumor stretches the surrounding gels circumferentially, with the maximum stress occurring at the spheroid boundary. Increasing the gel concentration, reduces the magnitude of the circumferential stresses outside the tumor because even though *c_H_* increases, the smaller tumors displace the gel less. At equilibrium, there is a net volume loss in the tumor center, which is balanced by volume gain at the boundary (Figure 2C, solid curves).

To examine the effect of water efflux, we also fit the data without considering feedback from the elastic stress (*γ*_*A*_ = 0). In this case, the predicted radii (dashed curves in Figure 2A) also provide a good fit of the data, but the stress distributions and net volume growth rates are more heterogeneous when considering growth regulated by nutrient level alone (Figures 2B and 2C, dashed). Further, using the corrected Akaike information criterion (AICc) [30] suggests the model with feedback provides a better fit to the experiment (see Text S2). For the water-efflux feedback function (Eq.(9)), we have considered the effect of the circumferential stress 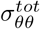, reasoning that it dominates the mechanical state of the cells because it represents stresses from two principal directions orthogonal to the radial direction. One could change 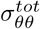 to the stress invariant 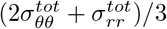 and obtain similar results. See Text S2 and Figure S3 in Text S2.

We calculate the average cell density in the tumor by 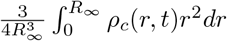 and plot this as a function of time. The average density increases with time. For more constrained tumors with increased gel stiffness, the average cell density increases, consistent with experimental data (vertical line segments in Figure 2D). Additionally, our simulation reveals that the region with largest cell density shifts towards the center of tumor (Figure 3A), indicating that tumor cells are most packed inside due to the inward cell flow. In this case, the effect of cell flux on the local cell density becomes primary, because volume growth and proliferative activities slow down due to globally elevated compression from the spatial confinement.

**Fig 3.**
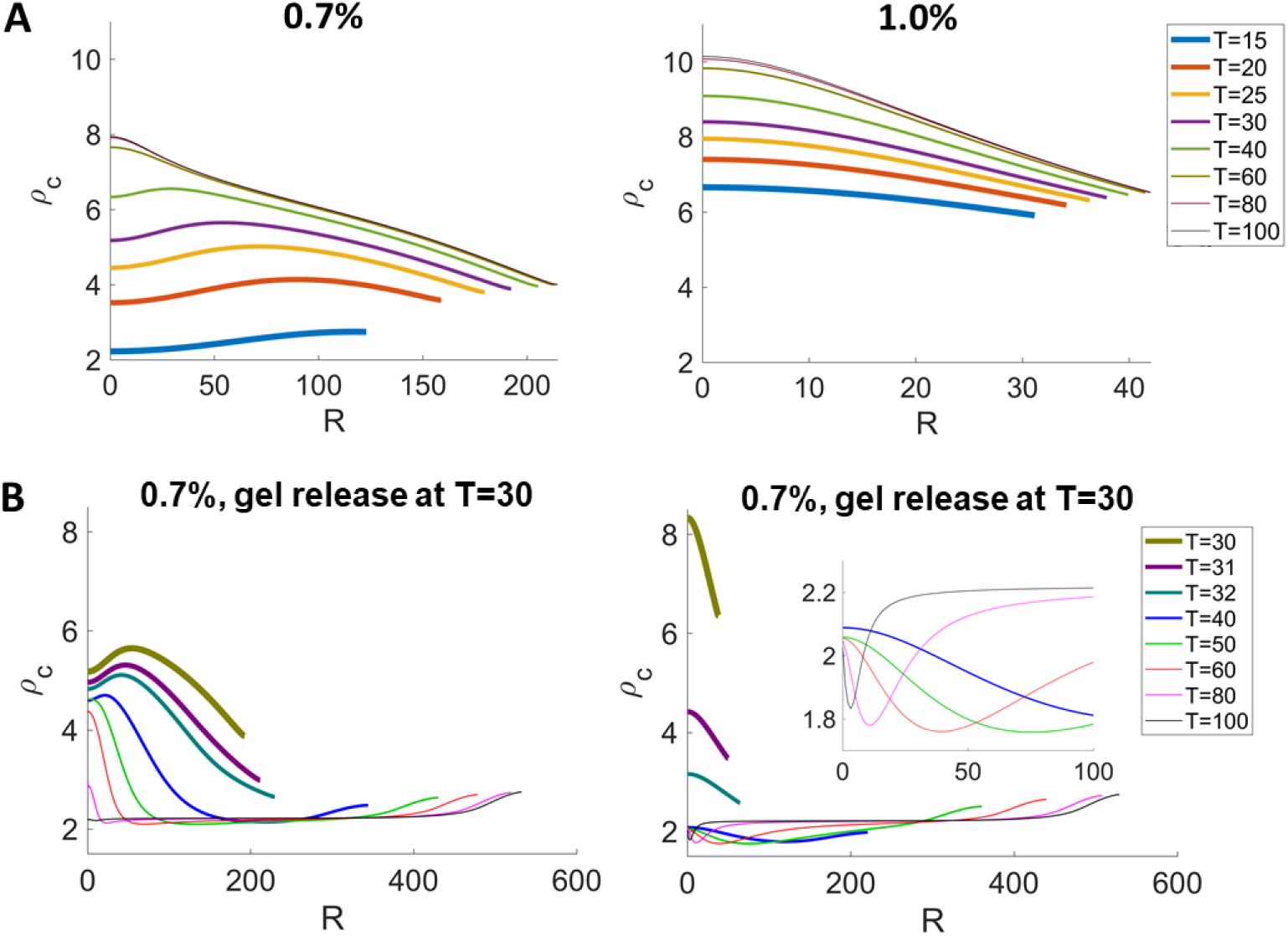
Cell densities *ρ_c_* in the constrained tumor in Figure 2. (A) The regions with largest cell density shifts towards the tumor center, indicating cells are more packed inside the tumor. (B) After releasing the gel, the cell density recovers to the same level in the unconstrained tumor. See Text S3 for the list of parameters.

To model the gel-removal experiments from [10] (Figure 2E-F), we set *c_H_* = 0 in all the gel cases and use the common set of fitted tumor-associated parameters (see Text S2 for details). Again, there is good quantitative agreement between the numerics and experiments, which both tend to recover the growth of the unconstrained spheroid. The average cell density also recovers to the same level as in the free-boundary case. (Figure 2E and Figure 2F) In addition, we also show that the spatial distribution of the cell density reverse to the distribution in the free-boundary case upon gel-removal, where the cell density near the tumor boundary becomes higher again (compare Figure 3B with Figure 1E).

We also use the data from [12,13], where colon carcinoma tumor spheroids containing mouse CT26 cell lines where grown under isotropic compression from an osmotically-induced external pressure. As suggested by [12, 13], the compressive stresses reduce proliferation rates without increasing apoptosis. Therefore, we fit our model with feedback on the proliferation rate to data in Figure 1 in [13]. In this case, the tumor radius is determined by the sensitivity to feedback from mechanical stresses (γ_λ_) and pressure boundary condition 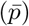, in addition to the volume loss rate λ_*A*_ and other parameters. We fit the experimental data in tumor radius (shown as symbols in Figure 4A) and data of relative cell density, where the density in the compressed tumor at tumor center is approximately 20% larger than that in the free tumor (see the triangle (mean) with error bars in Figure S1 in Text S4). Our fitting yields good quantitative agreement between the model and experiments for the dynamics of the tumor spheroid radii (Figure 4A), as well as the cell density (Figure S1 in Text S4). Similar to Figure 2B, the circumferential stress is also compressive at the tumor boundary, and becomes tensile towards the tumor center. Both the stress and the net volume growth are more uniform when feedback is considered. This can be seen by comparing the solid curve (with feedback) and dashed curve (without feedback) in Figure 4B-C.

**Fig 4.**
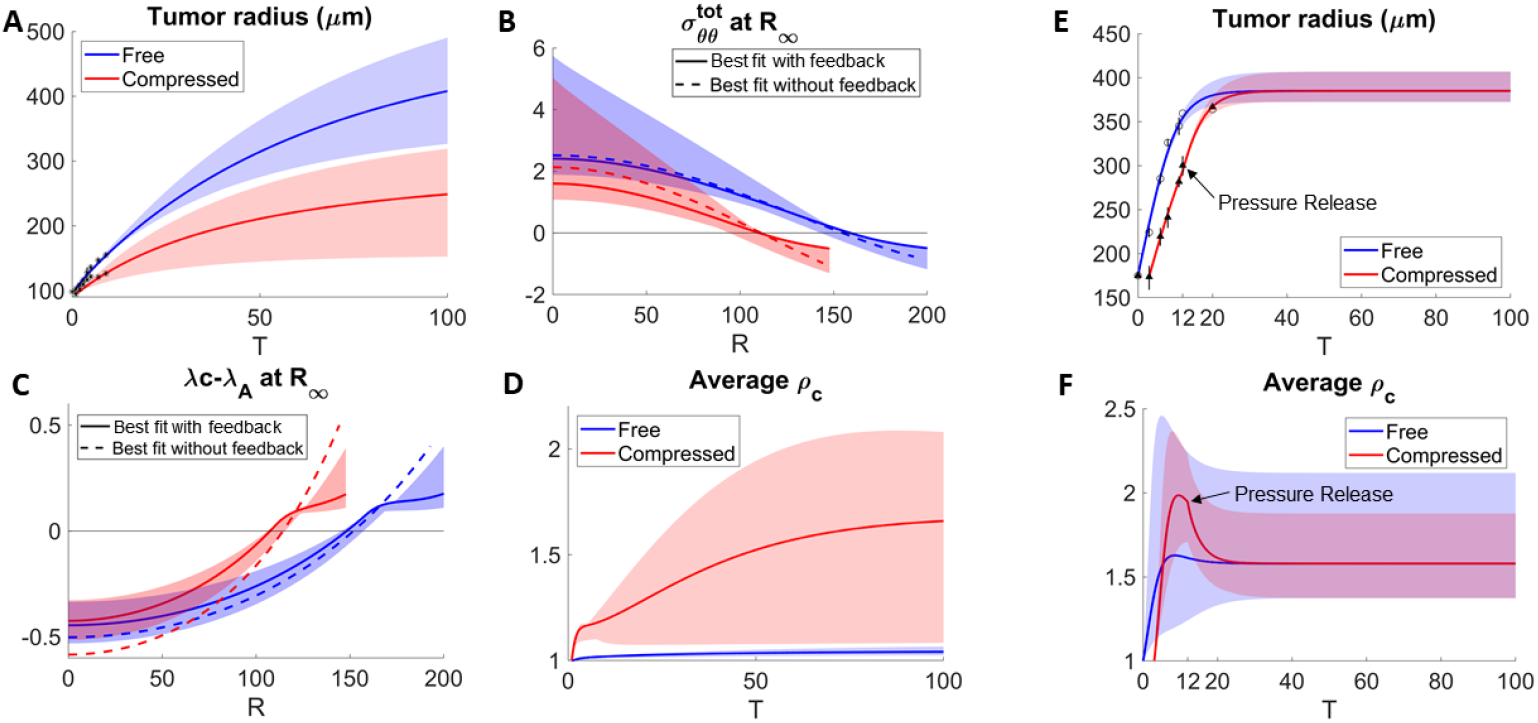
Fitting the model to data from [12, 13] of tumor spheroids grown free or with external pressure, with and without (dashed) feedback from the elastic stress (A). External pressure is released at the time reported in [13] (E). Bands show the results within 10% of the best fit with feedback in all panels. (B) and (C): The distributions of stresses and net volume growth rates from the model. (D) and (F): Average cell density of tumors in (A) and (E) respectively. See Text S4 for the list of parameters.

Consistent to the trend in Figure 2D, the average cell density is higher in the more compressed tumor (Figure 4D). Again, external extra compression shifts the region with higher cell density towards tumor center, as can be seen in Figure S1 in Text S4. The tumor radius, the average cell density, as well as the cell density distribution are all reversible upon the pressure removal (Figure 4E, 4F, and Figure S1 in Text S4).

### Tumor size, stress and density patterns at equilibrium

Next, we perform a parametric study to investigate how the internal stress relaxation and external spatial confinement influence the sizes of tumor spheroids, their stress distributions and anisotropies. At equilibrium, tumor sizes decrease with *c_H_* and increase with *β* due to external loading and internal relaxation, respectively (Figure 5A). The white dashed curve marks the boundary between tumor spheroids with tensile (to the left) and compressive (to the right) stresses at the spheroid center. When both *c_H_* and *β* increase, the stress is less elastic and more hydrostatic (Figure 5B, spatial distributions in Figure S1-S2 in Text S5) and less anisotropic (Figure 5C, spatial distributions in Figure S3-S4 in Text S5). In addition, the stress distributions along the tumor radii (Figure 5D) become more uniform and less sensitive to changes in *β* when *c_H_* increases, since the stress are dominated by compression from the external loading. When *β* and *c_H_* lie within the region marked by the dashed red curve in Figure 5A, the tumors are small but their stress distributions can be heterogeneous (Figure 5D). Interestingly, the corresponding dynamics of the spheroid radii are non-monotone as the stress equilibrates slowly towards the steady-state (Figure S5 in Text S5); we have not seen this in published data, however. Following [31], we anticipate this could lead to a break in radial symmetry.

**Fig 5.**
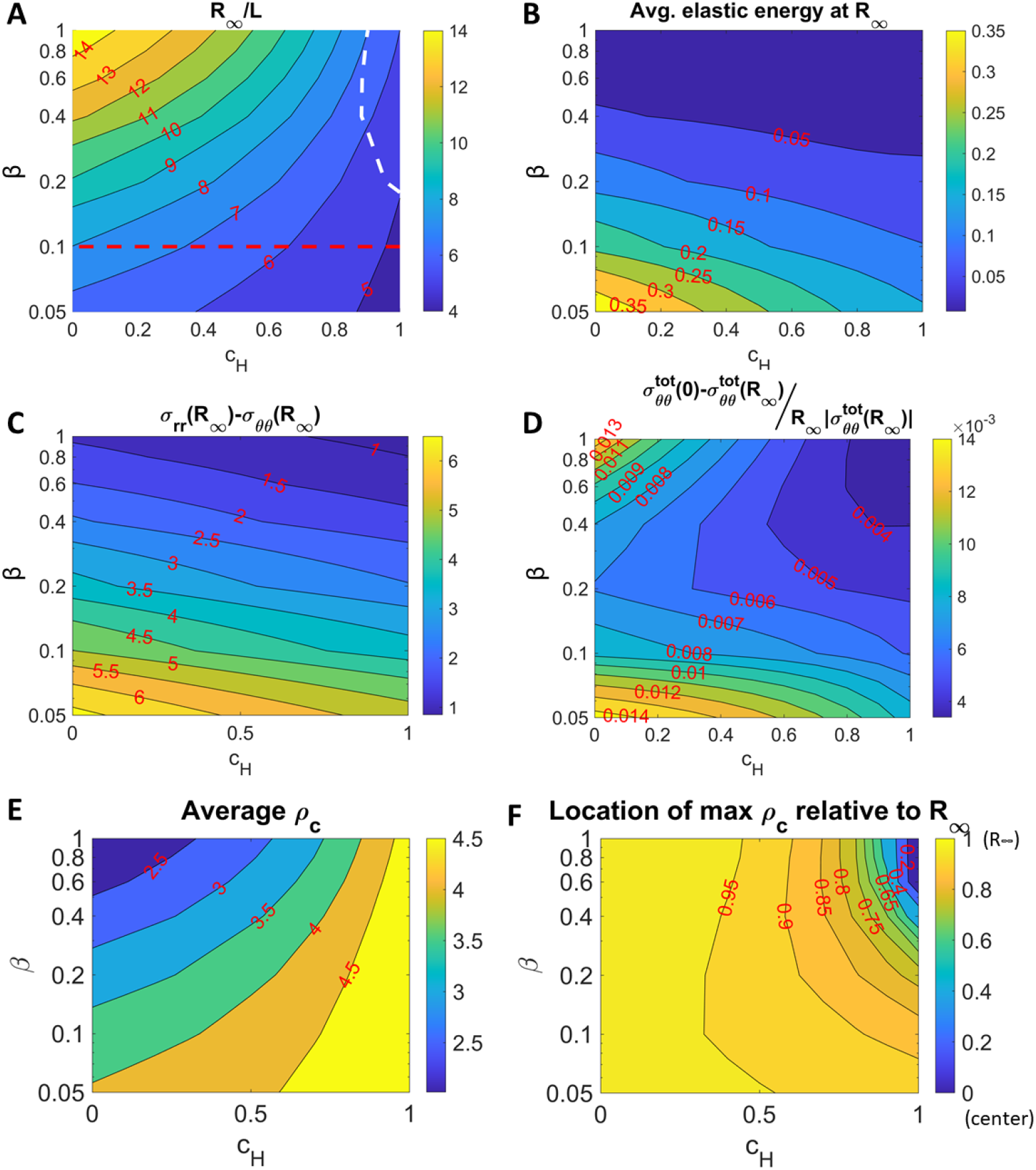
Contour plot of steady-state tumor radius (A), average elastic energy in the tumor at steady-state (B), stress anisotropy at tumor boundary (C), averaged stress gradient in the tumor, (D), steady-state average density (E), and location of maximum *ρ_c_* relative to tumor boundary (F) as a function of *β* and *c_H_*. The dashed white line shows the 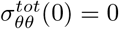 contour. The dashed red line shows the region where the dynamics of the radius are non-monotone (see Figure S1 in Text S5). In (F), a value of 1 corresponds to the tumor boundary, while a value of 0 corresponds to the tumor center. See the main text for more details.

We also investigate how the stress relaxation and external spatial confinement affects the magnitude and spatial distribution of cell density. At equilibrium, the average cell density increases with external confinement, but decreases with stress relaxation (Figure 5E). As the external confinement becomes stronger, cells are more packed near the tumor center (Figure 5F).

## Discussion

We considered the influence of mechanical stress on rates of tumor tissue volume changes via the the net effect between cell cycle in which the cell increases in size and the water efflux in which the cell decreases in size. The cell cycle can respond to the mechanical stresses via the shutting the transcriptional coactivators YAP and TAZ between the cytoplasm and the nucleus. The translocation of YAP and TAZ proteins is known to respond to multiple inputs, including the Hippo signaling pathway [32]. The water efflux has been suggested by [13] as the explanation of the tumor spheroid shrinkage in response to external compression. By considering both effects, we found good agreement with experimental data from [10, 12, 13] where the volumetric growth rates of tumor spheroids adjust as the level of external confinement [10] or hydrostatic compression [12,13] (presented in Text S4) varies. We have also modeled the spatial temporal dynamics of the cell density–the number of cells per volume–in the growing tumor where the local cell density is increased by the cell division rate, decreased (diluted) by the local volume growth, and further adjusted by the local cell flux to due advection or random neighbor exchanges. To prevent division of very small cells, we further consider that the division rate is decreased by local increases in cell density. Multiple feedback machineries such as contact inhibition [22] and cell cycle checkpoints [23] may be responsible for this negative feedback loop. By fitting the cell density data from [10,13] simultaneously with the data of the volumetric growth rates, we found not only that the average cell density increases with the strength of external compression or confinement, but also that the location with the maximal cell density transits from the tumor boundary to the center.

To model the spatial temporal dynamics of the stresses in the tumor tissue, we have developed and applied a new model of stress relaxation using an Eulerian framework. At the macroscopic tissue level, we have coupled the rate of growth with diffusion-limited nutrient transport, which results in differential accumulated growth. This is different from previous works that prescribe the differential accumulated growth as spatial varying functions [31, 33] and is facilitated by the full Eulerian framework. To relax the stress, we have introduced a relaxation rate *β* and the resulting system is similar to but simpler than the classical Maxwell model of viscoelasticity. The cytoskeletal and intercellular junction remodeling should affect *β*. For example, faster turnover/reassembly of these structural and adhesion molecules should increase *β*. By fitting spheroid data from these two independent studies using different tumor cell lines, we find the relaxation rate *β* ~ 1 per day, which is comparable to the volumetric growth rate of the tumor. Although we have not explicitly modeled the extracellular matrix in the growing tumor, we note that the dynamics of matrix and cell-matrix interactions should also impact the stress accumulation and relaxation. This is left to the future work. Our model predicts that feedback from elastic stresses result in a more uniform spatial pattern of growth rates, which is analogous to the spatial patterning of cell proliferation observed during development of the *Drosophila* wing disc [19–21] where feedback from elastic stresses was also found to be important for this pattern of growth. We found that when the compression from external confinement is non-negligible compared to the internal compression generated by differential growth, the tumor spheroid sizes and the stress distributions are not sensitive to changes in the material properties of the spheroids. Further, the total stress is nearly uniform and is dominated by hydrostatic pressure.

We can also gain insight on tumor growth *in vivo*. When a tumor increases its instantaneous elasticity relative to the external confinement (e.g., decreasing *c_H_*), such tumors are more able to change their size and mechanical state autonomously. This may explain why increased tumor stiffness is an established hallmark in tumor malignancy [18,34]. On one hand, when *c_H_* is small, the tumor size as well as the average cell size (the inverse of cell density) can be increased by increasing the stress relaxation rate *β*. On the other hand when *β* is small, the tumors have large elastic energy and anisotropy and the model predicts that the dynamics of the tumor can be non-monotone, although we have not seen this observed in published data. However, we anticipate that such tumors may be subject to morphological instability, increasing local invasiveness. In summary, we suggest that when the tumor stiffness dominates over the surrounding compressive stresses, then active relaxation – an effect from lumping the turnover and deposition of intracellular cytoskeleton structures, intercellular adhesion complexes, and extracellular matrices–can be used to leverage against local invasiveness and bulk expansion during tumor progression.

## Conclusion

We have developed biomechanical model that accounts for the stress generation and relaxation in the growing tumor spheroid, and have considered the chemomechanical responses of tumor tissue in regulating the size distribution of cells and the integrated size of the tissue. By fitting the model to experimental data from two independent studies of tumor spheroid growth and their cell density distributions, treating the tumors as incompressible, neo-Hookean elastic materials, we find that the rates of stress relaxation of tumor tissues can be comparable to volumetric growth rates. Our study provides insight on how the biophysical properties of the tumor and host microenvironment, mechanical feedback control and diffusion-limited differential growth act in concert to regulate spatial patterns of stress and growth. In particular, when the tumor is stiffer than the host, our model predicts tumors are more able to change their size and mechanical state autonomously, which may help to explain why increased tumor stiffness is an established hallmark of malignant tumors.

## Supporting information

Supplemental Data for Figure 1

Supplemental Data for Figure 2

Supplemental Data for Figure 3

Supplemental Data for Figure 4

Supplemental Data for Figure 5

## Supporting information

**S1 Text Supplemental data for Figure 1.** Growth-induced deformation in an Eulerian frame, non-dimensionalization, radially symmetric case, numerical methods, parameters and supporting figures for Figure 1.

**S2 Text Supplemental data for Figure 2.** Grid-search parameter optimization, parameters and supporting figures for Figure 2.

**S3 Text Supplemental data for Figure 3.** Parameters for Figure 3.

**S4 Text Supplemental data for Figure 4.** Grid-search parameter optimization, parameters and supporting figures for Figure 4.

**S5 Text Supplemental data for Figure 5.** Parameters and supporting figures for Figure 5.

## Acknowledgments

J.L., H.Y. and D.R-G. acknowledge partial funding from the National Science Foundation-Division of Mathematical Sciences (NSF-DMS) grants DMS-1714973 and DMS-1763272/Simons Foundation (594598, QN) for the Center for Multiscale Cell Fate Research at UC Irvine. M.W acknowledge partial funding from NSF-DMS-2012330. JL additionally acknowledges partial funding from the National Institutes of Health (NIH) grant 1U54CA217378-01A1 for a National Center in Cancer Systems Biology at the UC Irvine (also provided support to H.Y.), and from NIH grant P30CA062203 for the Chao Family Comprehensive Cancer Center at UC Irvine.

