## Supplemental Data for Figure 1 for "Stress generation, relaxation and size control in confined tumor growth"

(Dated: August 29, 2021)

### 1. SUPPLEMENTAL DATA FOR FIGURE 1

#### 1.1. Growth-induced deformation in an Eulerian frame

##### *1.1.1. The constitutive relation: an overdamped nonlinear elastic system*

We assume the internal stress between cells and extracellular matrices balances the dissipative forces from the other biomaterials such as interstitial fluids, blood vessels, cell debris, etc. For simplicity, we consider the constitutive law

$$\overline{\alpha}\overline{\mathbf{v}}(\overline{\mathbf{x}},\overline{t}) = \overline{\nabla}\cdot\overline{\tilde{\boldsymbol{\sigma}}}(\overline{\mathbf{x}},\overline{t}) = \overline{\nabla}\cdot\overline{\boldsymbol{\sigma}}(\overline{\mathbf{x}},\overline{t}) - \overline{\nabla}\overline{p}(\overline{\mathbf{x}},\overline{t}), \tag{1}$$

where $\overline{\mathbf{v}}$ is the velocity, $\overline{\alpha}$ the friction (or drag) coefficient, and the internal stress (the Cauchy stress tensor) is $\overline{\tilde{\boldsymbol{\sigma}}}(\overline{\mathbf{x}},\overline{t}) = \overline{\boldsymbol{\sigma}}(\overline{\mathbf{x}},\overline{t}) - \overline{p}(\overline{\mathbf{x}},\overline{t})\mathbf{I}$ where $\overline{p}$ is the pressure, and $\overline{\nabla}$ is the gradient in $\overline{\mathbf{x}}$, which is a spatial point in physical space (current configuration). The Cauchy stress tensor $\overline{\tilde{\boldsymbol{\sigma}}}(\overline{\mathbf{x}},\overline{t})$ will be derived from a hyperelastic energy function and the pressure $\overline{p}(\overline{\mathbf{x}},\overline{t})$ will ensure incompressibility. The derivation can also be extended to more general nonlinear elastic energies [1]. Finite, or nonlinear, elasticity is traditionally posed in a Lagrangian frame [2, 3], e.g., stress is obtained at a point $\overline{\mathbf{X}}$ in the reference configuration. The overbars denote dimensional variables. In the following, we derive the viscoelastic elastic tumor model in the Eulerian frame (current configuration) presented in the main text. Here, we do not assume radial symmetry and elastic stresses are generated by accumulated finite volumetric growth.

##### *1.1.2. Growth-induced stress in finite elasticity*

**Mass balance and incompressibility**

Given the local net volumetric growth rate $\Gamma(\overline{\mathbf{x}},\overline{t})$, mass conservation gives

$$\frac{\partial\overline{\rho}(\overline{\mathbf{x}},\overline{t})}{\partial\overline{t}} + \overline{\nabla}\cdot(\overline{\rho}(\overline{\mathbf{x}},\overline{t})\overline{\mathbf{v}}(\overline{\mathbf{x}},\overline{t})) = \overline{\rho}(\overline{\mathbf{x}},\overline{t})\overline{\Gamma}(\overline{\mathbf{x}},\overline{t}), \tag{2}$$

where $\overline{\rho}(\overline{\mathbf{x}},\overline{t})$ is the cell density field. Assuming the tissue is strictly incompressible ($\overline{\rho}(\overline{\mathbf{x}},\overline{t}) = \text{const.}$), we have

$$\overline{\nabla}\cdot\overline{\mathbf{v}} = \overline{\Gamma}(\overline{\mathbf{x}},\overline{t}). \tag{3}$$

**Accumulated volume variation and growth tensors**

The accumulated volume variation $J$ and the rate of growth $\overline{\Gamma}$ are related via

$$\frac{\partial J(\overline{\mathbf{x}},\overline{t})}{\partial\overline{t}} + \overline{\mathbf{v}}(\overline{\mathbf{x}},\overline{t})\cdot\overline{\nabla}J(\overline{\mathbf{x}},\overline{t}) = J(\overline{\mathbf{x}},\overline{t})\overline{\Gamma}(\overline{\mathbf{x}},\overline{t}). \tag{4}$$

Following [4], we define the growth tensor to be a deformation tensor $\mathbf{G}(\overline{\mathbf{X}},\overline{t})$ from the initial configuration to the unstressed grown configuration. See Figure 1, $A \to B1$. Here, we assume $\mathbf{G}(\overline{\mathbf{X}},\overline{t})$ is a symmetric stretch tensor:

$$\mathbf{G} = \Sigma_{i=1}^{3}\lambda_i\hat{\mathbf{n}}_i\otimes\hat{\mathbf{n}}_i, \tag{5}$$

where $\lambda_i$'s are the principal growth magnitudes and $\hat{\mathbf{n}}_i$'s are the principal growth directions. To be consistent with volumetric variation the relation

$$\det(\mathbf{G}) = \Pi_{i=1}^{3}\lambda_i = J \tag{6}$$

needs to be satisfied. So, when the growth is isotropic ($\lambda_1 = \lambda_2 = \lambda_3$), the growth tensor is

$$\mathbf{G} = J^{1/3}\mathbf{I}, \tag{7}$$

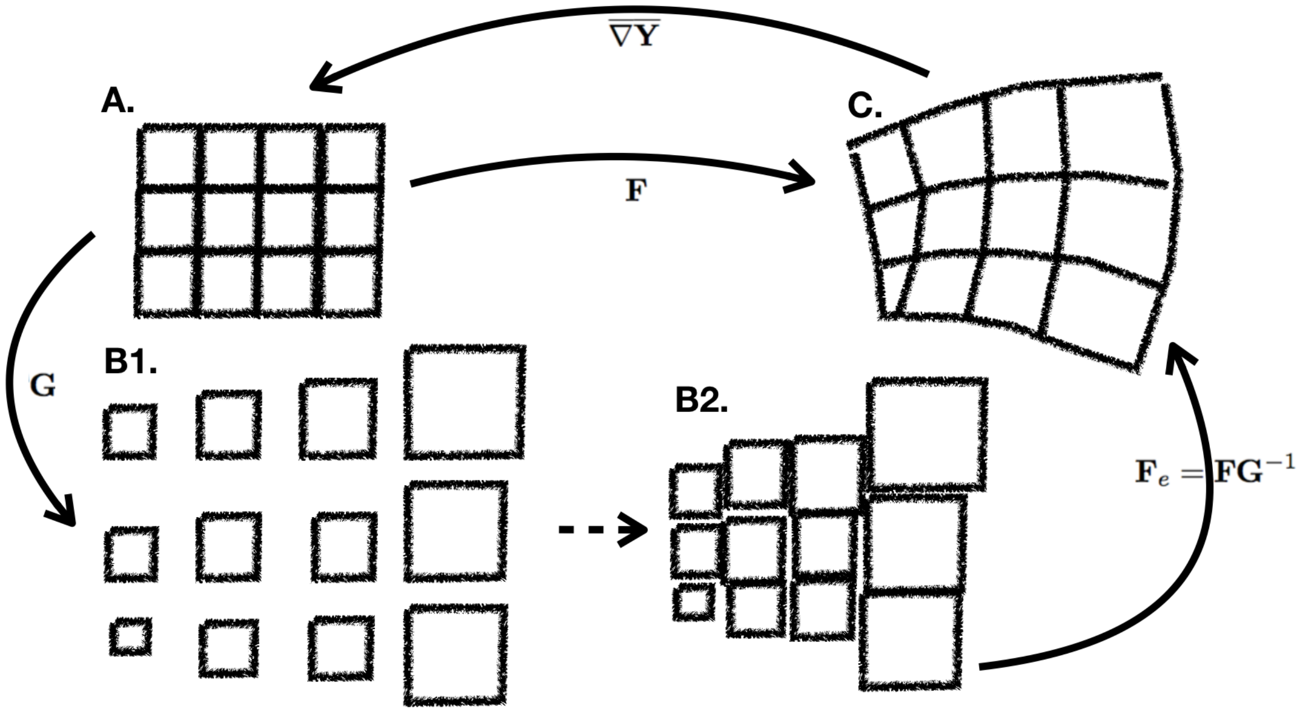


Fig.S 1: A. Configuration before growth; B1. Configuration after each box of material grows freely; B2. The incompatible configuration that arises from putting the boxes together after growth; C. Configuration after isochoric deformations, which results in internal stresses.

where $\mathbf{I}$ is the identity tensor.

#### Growth incompatibility and finite elastic deformation

At time $t$, $\mathbf{G}$ operating on the material $\overline{\mathbf{X}}$ may result in incompatible configurations (Fig. 1, $A \to B1 \to B2$) and the blocks in the continuum body have to be locally stretched and rotated (Fig. 1, $B2 \to C$), through elastic deformation, in order to remain connected and non-overlapping in the current configuration (physical space). We denote the elastic deformation by a tensor-valued function $\mathbf{F}_e$, which is related to the total geometric deformation tensor $\mathbf{F}$ (Fig. 1, $A \to C$) and the growth tensor $\mathbf{G}$ by [4]

$$\mathbf{F}_e = \mathbf{F}\mathbf{G}^{-1}. \tag{8}$$

In other words, the stress-free reference configuration is not the initial configuration, but a virtual configuration after growth. Although $J_e = \det(\mathbf{F}_e) = 1$ due to incompressibility, the components of $\mathbf{F}_e$ may be far from unity, which necessitates the use of finite (nonlinear) elasticity.

#### Stress in finite elasticity with incompressibility

We derive the stresses using nonlinear elasticity theory [2, 3]. Here we assume that the tissues during growth behave as a hyperelastic material and are able to sustain finite deformations without material failure. Given a strain energy function $W(\mathbf{F}_e)$, representing the energy per reference volume (Joules per unit volume), the Cauchy stress tensor can be derived by the virtual displacement principle:

$$\bar{\tilde{\boldsymbol{\sigma}}} = J_e^{-1} \frac{\partial W}{\partial \mathbf{F}_e} \mathbf{F}_e^T. \tag{9}$$

This formula is generally true when $J_e = \det(\mathbf{F}_e) = 1$ is not imposed. Assuming the tissue to be isotropic, the strain energy function can be rewritten as $W(\mathbf{F}_e) = W(I_1, I_2, I_3)$, where $I_1 = \lambda_1^2 + \lambda_2^2 + \lambda_3^2$, $I_2 = \lambda_1^2\lambda_2^2 + \lambda_1^2\lambda_3^2 + \lambda_2^2\lambda_3^2$, and $I_3 = \lambda_1^2\lambda_2^2\lambda_3^2$ and the $\lambda_i$'s are the eigenvalues of $[\mathbf{F}_e\mathbf{F}_e^T]^{1/2}$ (or $[\mathbf{F}_e^T\mathbf{F}_e]^{1/2}$) and $I_1$, $I_2$, and $I_3$ are the principal invariants of $[\mathbf{F}_e\mathbf{F}_e^T]^{1/2}$ (or $[\mathbf{F}_e^T\mathbf{F}_e]^{1/2}$). Using a change of variables and the chain rule, the Cauchy stress tensor can be rewritten as

$$\bar{\tilde{\boldsymbol{\sigma}}} = 2J_e^{-1}\left((I_2\frac{\partial W}{\partial I_2} + I_3\frac{\partial W}{\partial I_3})\mathbf{I} + \frac{\partial W}{\partial I_1}\mathbf{F}_e\mathbf{F}_e^T - I_3\frac{\partial W}{\partial I_2}[\mathbf{F}_e\mathbf{F}_e^T]^{-1}\right). \tag{10}$$

Since we assume incompressibility, we can derive $I_3 = \det(\mathbf{F}_e)^2 = 1$, and so we need to drop the term $I_3\frac{\partial W}{\partial I_3}$ and introduce the pressure field $\bar{p}$ which serves as a Lagrangian multiplier for the constraint $I_3 = 1$. By assuming $\frac{\partial W}{\partial I_1} = const.$ and $I_3\frac{\partial W}{\partial I_2} = const.$ and absorbing $I_2\frac{\partial W}{\partial I_2}$ into the pressure field, we obtain the incompressible Mooney-Rivlin model for the Cauchy stress tensor:

$$\bar{\tilde{\boldsymbol{\sigma}}} = \bar{\mu}_1\mathbf{F}_e\mathbf{F}_e^T + \bar{\mu}_2[\mathbf{F}_e\mathbf{F}_e^T]^{-1} - \bar{p}\mathbf{I}, \tag{11}$$

where $\bar{\mu}_1$ and $\bar{\mu}_2$ are elastic coefficients. Since the tissue is incompressible and growth is isotropic, we have $\mathbf{G} = J^{1/3}\mathbf{I} = \det(\mathbf{F})^{1/3}\mathbf{I}$. Therefore, the Cauchy stress tensor becomes

$$\bar{\tilde{\boldsymbol{\sigma}}} = \bar{\mu}_1 J^{-2/3}\mathbf{F}\mathbf{F}^T + \bar{\mu}_2 J^{2/3}[\mathbf{F}\mathbf{F}^T]^{-1} - \bar{p}\mathbf{I}. \tag{12}$$

#### 1.1.3. *Reformulating stresses using reference map*

In order to rewrite the stress $\bar{\tilde{\boldsymbol{\sigma}}}$ in physical coordinates (current configuration), we introduce a reference map $\overline{\mathbf{Y}}$ that relates the current and reference configurations, following [5, 6]. In particular, if the initial configuration is stress-free, then $\overline{\mathbf{Y}}$ captures the point $\overline{\mathbf{X}}$ at $t = 0$ (reference configuration) where the material at position $\bar{\mathbf{x}}$ at time $\bar{t}$ came from. That is, $\bar{\mathbf{Y}}(\bar{\mathbf{x}}, t) = \bar{\mathbf{X}}$. The reference map evolves via

$$\frac{\partial\overline{\mathbf{Y}}}{\partial\bar{t}} + \bar{\mathbf{v}}\cdot\overline{\nabla}\overline{\mathbf{Y}} = \bar{0}, \quad \overline{\mathbf{Y}}(\bar{\mathbf{x}}, \bar{0}) = \bar{\mathbf{x}} = \overline{\mathbf{X}}. \tag{13}$$

The total deformation is given by $\mathbf{F} = [\overline{\nabla}\overline{\mathbf{Y}}]^{-1}$. Putting everything together we obtain

$$\bar{\tilde{\boldsymbol{\sigma}}}(\bar{\mathbf{x}}, \bar{t}) = \bar{\mu}_1 J^{-2/3}[\overline{\nabla}\overline{\mathbf{Y}}]^{-1}[\overline{\nabla}\overline{\mathbf{Y}}]^{-1T} + \bar{\mu}_2 J^{2/3}[\overline{\nabla}\overline{\mathbf{Y}}]^T[\overline{\nabla}\overline{\mathbf{Y}}] - \bar{p}\mathbf{I} \tag{14}$$

where $J = \det(\mathbf{F}) = [\det(\overline{\nabla}\overline{\mathbf{Y}})]^{-1}$. Notice the stress formula above corresponds to a neo-Hookean incompressible material when $\bar{\mu}_2 = 0$. From now on, we assume $\bar{\mu}_2 = 0$. With $\alpha = 0$ in Eq. (1), the system of Eqs. (1),(3),(4),(13) and (14) is equivalent to the growth elasticity theory when the tissue is assumed to be an incompressible neo-Hookean material and the growth is isotropic (see [7–9]). In previous models, instead of solving Eq. (4) for the accumulated growth $J$ from $\overline{\Gamma}(\bar{\mathbf{x}}, \bar{t})$, $J$ is prescribed as a function of space either in the Lagrangian or Eulerian frame (see the isotropic growth case when $\gamma_1 = \gamma_2 = const.$ in [7] and $g(r) = 1 + \nu(r - a)$ in [8]). We reason that even when the accumulated growth $J$ is defined as a function of the current coordinate $J(\bar{\mathbf{x}})$, as in [8], it is essentially a Langrangian formulation, which can be seen by the change of variables $\tilde{J}(\overline{\mathbf{X}}) = J(\bar{\mathbf{x}}(\overline{\mathbf{X}}, t))$. This means that in this previous framework [8], the accumulated growth only depends on the initial position and the deformation state of the tissue, which limits its dependence on other factors such as the chemical signals. Our formulation allows a straightforward coupling between the growth rate $\overline{\Gamma}(\bar{\mathbf{x}}, \bar{t})$ and biochemical signals (e.g., nutrient, growth factors, and oxygen), and the spatial distribution of accumulated growth is no longer requires simplifying assumptions. We note that the Lagrangian frame makes it difficult to couple growth with biochemical fields, which are naturally posed in the Eulerian frame (e.g., reaction and diffusion in the current configuration) [10]. By introducing the evolution of the reference map in Eq.(13), we are able to describe the coupling among reaction-diffusion processes of chemical signals, growth and mechanics fully in the same Eulerian frame.

As a result of Eq.(13), we can derive the evolution of the stress $\bar{\tilde{\boldsymbol{\sigma}}}(\bar{\mathbf{x}}, \bar{t})$ in the following manner. By taking the spatial gradient of Eq. (13) in physical coordinates, one can first derive

$$\frac{\partial[\overline{\nabla}\overline{\mathbf{Y}}]}{\partial\bar{t}} + \mathbf{v}\cdot\overline{\nabla}[\overline{\nabla}\overline{\mathbf{Y}}] \;=\; -[\overline{\nabla}\overline{\mathbf{Y}}]\overline{\nabla}\mathbf{v}, \tag{15}$$

and its transpose

$$\frac{\partial[\overline{\nabla}\overline{\mathbf{Y}}]^T}{\partial\bar{t}} + \mathbf{v}\cdot\overline{\nabla}[\overline{\nabla}\overline{\mathbf{Y}}]^T \;=\; -\overline{\nabla}\mathbf{v}^T[\overline{\nabla}\overline{\mathbf{Y}}]^T. \tag{16}$$

Then, using $\mathbf{I} = [\overline{\nabla}\mathbf{Y}][\overline{\nabla}\mathbf{Y}]^{-1} = [\overline{\nabla}\mathbf{Y}]^T[\overline{\nabla}\mathbf{Y}]^{-1T}$ and $\frac{D\mathbf{I}}{D\bar{t}} = \frac{\partial \mathbf{I}}{\partial \bar{t}} + \overline{\mathbf{v}}\cdot\overline{\nabla}\mathbf{I} = 0$, one can derive the convection equations for $[\overline{\nabla}\mathbf{Y}]^{-1}$ and $[\overline{\nabla}\mathbf{Y}]^{-1T}$ from Eqs. (15) and (16), respectively:

$$\frac{\partial[\overline{\nabla}\mathbf{Y}]^{-1}}{\partial \bar{t}} + \mathbf{v}\cdot\overline{\nabla}[\overline{\nabla}\mathbf{Y}]^{-1} \;=\; \overline{\nabla}\mathbf{v}[\overline{\nabla}\mathbf{Y}]^{-1} \tag{17}$$

$$\frac{\partial[\overline{\nabla}\mathbf{Y}]^{-1T}}{\partial \bar{t}} + \mathbf{v}\cdot\overline{\nabla}[\overline{\nabla}\mathbf{Y}]^{-1T} \;=\; [\overline{\nabla}\mathbf{Y}]^{-1T}\overline{\nabla}\mathbf{v}^T. \tag{18}$$

Using $\frac{D(\det \mathbf{A})}{Dt} = \frac{\partial(\det \mathbf{A})}{\partial \mathbf{A}} : \frac{D\mathbf{A}}{Dt} = (\det \mathbf{A})\mathrm{tr}(\mathbf{A}^{-1}\frac{D\mathbf{A}}{Dt})$, where $\frac{D}{Dt} = \partial/\partial_t + \mathbf{v}\cdot\nabla$, one can derive

$$\frac{D(\det[\overline{\nabla}\mathbf{Y}])}{D\bar{t}} \;=\; -\det[\overline{\nabla}\mathbf{Y}]\overline{\nabla}\cdot\mathbf{v} \tag{19}$$

$$\frac{D(\det[\overline{\nabla}\mathbf{Y}^{-1}])}{D\bar{t}} \;=\; \det[\overline{\nabla}\mathbf{Y}^{-1}]\overline{\nabla}\cdot\mathbf{v} \tag{20}$$

from Eqs. (15) and (17), respectively. Notice Eq.(20) is equivalent to Eq.(4) since $J = \det(\mathbf{F}) = [\det(\overline{\nabla}\mathbf{Y}^{-1})]$ and $\overline{\nabla}\cdot\mathbf{v} = \Gamma$. This means the accumulated volume growth is automatically tracked by the dynamics of $\overline{\mathbf{Y}}$ from Eq. (13). For the non-pressure part of the stress

$$\overline{\boldsymbol{\sigma}}_e := \overline{\mu}_1[\det(\overline{\nabla}\mathbf{Y})]^{2/3}[\overline{\nabla}\mathbf{Y}]^{-1}[\overline{\nabla}\mathbf{Y}]^{-1T}, \tag{21}$$

we can derive its evolution as

$$\overset{\triangledown}{\overline{\boldsymbol{\sigma}}}_e := \frac{D\overline{\boldsymbol{\sigma}}_e}{D\bar{t}} - \overline{\nabla}\overline{\mathbf{v}}\;\overline{\boldsymbol{\sigma}}_e - \overline{\boldsymbol{\sigma}}_e\;\overline{\nabla}\overline{\mathbf{v}}^T = -\frac{2}{3}\,\overline{\Gamma}\overline{\boldsymbol{\sigma}}_e. \tag{22}$$

by using Eqs. (3), (17),(18), and (19). Given $\overline{\boldsymbol{\sigma}}_e$ and the velocity, the total Cauchy stress tensor $\overline{\tilde{\boldsymbol{\sigma}}}(\overline{\mathbf{x}}, \bar{t}) = \overline{\boldsymbol{\sigma}}_e - \bar{p}\mathbf{I}$ can be obtained by solving for the pressure from Eqs. (1) and (3). Although Eq. (22) holds, in fact $\overline{\boldsymbol{\sigma}}_e$ is solely determined from the reference map via Eq. (21).

If the tumor spheroid reaches an equilibrium size due to a balance between cell proliferation at the tumor periphery (volume gain) and cell apoptosis at the tumor core (volume loss), there is no mechanism from the above equation for the stress $\overline{\boldsymbol{\sigma}}_e$ to relax. This leads to a blow up of the stress in time (see main text -Results *A tumor spheroid growing with stress relaxation*), which has also been shown in a previous linear model considering isotropic growth [11]. The stress can be relaxed by introducing growth anisotropy [12] or viscoelasticity as noted by [13]. In the following, we introduce stress relaxation by modifying the dynamics of the reference map Eq. (13). We will demonstrate the corresponding stress relaxation behaves like a Maxwell viscoelastic model. Remarkably, our model formulation avoids solving the tensor equation that involves both the stress tensor and its upper-convected time derivative, thus allows more efficient computation.

#### 1.1.4. *Stress relaxation*

To model stress relaxation, we assume that reference configuration adapts to the current configuration at a rate $\overline{\beta}$. In the Eulerian frame, this can be modeled as:

$$\frac{\partial\overline{\mathbf{Y}}}{\partial \bar{t}} + \overline{\mathbf{v}}\cdot\overline{\nabla}\mathbf{Y} = \overline{\beta}(\mathbf{R}^T\overline{\mathbf{x}} - \overline{\mathbf{Y}}) \tag{23}$$

where $\mathbf{R}$ is the rotational part of the deformation gradient $\mathbf{F}$. The combination $\mathbf{R}^T\overline{\mathbf{x}}$ is needed to ensure objectivity of the system. From the polar decomposition of $\mathbf{F} = \mathbf{R}\mathbf{U}$, where $\mathbf{U} = \sqrt{\mathbf{F}^T\mathbf{F}}$ is the symmetric positive-definite stretch tensor in the material coordinate, we can uniquely define $\mathbf{R}$ as the local rotation (orthogonal transformation $\mathbf{R}^T = \mathbf{R}^{-1}$) that transforms differential elements from the tangent space in the material (curvilinear) coordinate into the tangent space in the current (curvilinear) coordinate (e.g., [14]). Consequently, we have

$$\begin{aligned}\frac{\partial[\overline{\nabla}\mathbf{Y}]}{\partial \bar{t}} + \mathbf{v}\cdot\overline{\nabla}[\overline{\nabla}\mathbf{Y}] \;&=\; -[\overline{\nabla}\mathbf{Y}]\overline{\nabla}\mathbf{v} + \overline{\beta}\left(\overline{\nabla}\left(\mathbf{R}^T\overline{\mathbf{x}}\right) - [\overline{\nabla}\mathbf{Y}]\right)\\ &=\; -[\overline{\nabla}\mathbf{Y}]\overline{\nabla}\mathbf{v} + \overline{\beta}(\mathbf{R}^T - [\overline{\nabla}\mathbf{Y}]) + \overline{\beta}\overline{\mathbf{x}}\cdot(\overline{\nabla}\mathbf{R})\end{aligned} \tag{24}$$

following the same differentiation procedure as in the previous section 1.1.3. Using $\frac{D\mathbf{A}^{-1}}{Dt} = -\mathbf{A}^{-1}\frac{D\mathbf{A}}{Dt}\mathbf{A}^{-1}$, we derive the evolution of the deformation gradient $\mathbf{F} = [\overline{\nabla}\overline{\mathbf{Y}}]^{-1}$ and obtain

$$\overset{\diamond}{\mathbf{F}} := \frac{\partial \mathbf{F}}{\partial \overline{t}} + \overline{\mathbf{v}} \cdot \overline{\nabla}\mathbf{F} - \overline{\nabla}\overline{\mathbf{v}} \cdot \mathbf{F} = -\overline{\beta}\mathbf{F}(\mathbf{R}^T - \mathbf{F}^{-1} + \overline{\mathbf{x}} \cdot (\overline{\nabla}\mathbf{R}))\mathbf{F} = \overline{\beta}\mathbf{F}(\mathbf{I} - \mathbf{U}) - \overline{\beta}\mathbf{F}(\overline{\mathbf{x}} \cdot (\overline{\nabla}\mathbf{R}))\mathbf{F}. \tag{25}$$

Notice that this equation is objective under the rotation $\overline{\mathbf{x}}^+ = \mathbf{Q}(t)\overline{\mathbf{x}}$ because on the left hand side we have $\overset{\diamond +}{\mathbf{F}^+} = \mathbf{Q}\overset{\diamond}{\mathbf{F}} = [\overset{\diamond}{\mathbf{F}}]^+$, and on the right hand side, the first term becomes $\mathbf{F}^+(\mathbf{I} - \mathbf{U}) = \mathbf{QF}(\mathbf{I} - \mathbf{U}) = [\mathbf{F}(\mathbf{I} - \mathbf{U})]^+$ and the second term becomes $\mathbf{F}^+(\overline{\mathbf{x}}^+ \cdot \overline{\nabla}_{\overline{\mathbf{x}}^+}\mathbf{R}^+)\mathbf{F}^+ = \left[\mathbf{QF}(\overline{\mathbf{x}} \cdot \overline{\nabla}_{\overline{\mathbf{x}}}\mathbf{R})\mathbf{F}\right] = [\mathbf{F}(\overline{\mathbf{x}} \cdot \overline{\nabla}_{\overline{\mathbf{x}}}\mathbf{R})\mathbf{F}]^+$, where we have omitted the $\overline{\beta}$ for simplicity.

In tumor spheroid growth, or in other symmetric growth configurations where no rotation is involved (e.g., $\mathbf{R} = \mathbf{I}$ and $\mathbf{F} = \mathbf{U}$), the right hand sides of Eqs. (23) and (25) become $\overline{\beta}(\overline{\mathbf{x}} - \overline{\mathbf{Y}})$ and $\overline{\beta}\mathbf{U}(\mathbf{I} - \mathbf{U})$, respectively. Note that the latter term can be interpreted as a nonlinear effect of the local stretches along the three (unchanged) principal directions.

One can further derive the evolution of the Finger deformation tensor $\mathbf{C} := \mathbf{FF}^T$ as

$$\overset{\triangledown}{\mathbf{C}} = \frac{\partial \mathbf{C}}{\partial \overline{t}} + \overline{\mathbf{v}} \cdot \overline{\nabla}\mathbf{C} - \overline{\nabla}\overline{\mathbf{v}} \cdot \mathbf{C} - \mathbf{C} \cdot \overline{\nabla}\overline{\mathbf{v}}^T = \overline{\beta}\left(\mathbf{R}\left(\mathbf{I} - \mathbf{U}\right)\mathbf{R}^T\mathbf{C} + \mathbf{CR}\left(\mathbf{I} - \mathbf{U}\right)\mathbf{R}^T\right) - \overline{\beta}\left(\mathbf{F}\left(\overline{\mathbf{x}} \cdot \overline{\nabla}\mathbf{R}\right)\mathbf{C} + \mathbf{C}\left(\overline{\mathbf{x}} \cdot \overline{\nabla}\mathbf{R}\right)^T\mathbf{F}^T\right). \tag{26}$$

Using $\frac{D(\det \mathbf{A})}{Dt} = \frac{\partial(\det \mathbf{A})}{\partial \mathbf{A}} : \frac{D\mathbf{A}}{Dt} = (\det \mathbf{A})\mathrm{tr}(\mathbf{A}^{-1}\frac{D\mathbf{A}}{Dt})$, we get the evolution of the "effective" accumulated volume variation $J = \det \mathbf{F}$ tracked by the adaptive reference map:

$$\frac{\partial J}{\partial \overline{t}} + \overline{\mathbf{v}} \cdot \overline{\nabla}J = \overline{\Gamma}J + \overline{\beta}\left(\mathrm{tr}(\mathbf{I} - \mathbf{U}) - \mathrm{tr}\left(\mathbf{F}\left(\overline{\mathbf{x}} \cdot \overline{\nabla}\mathbf{R}\right)\right)\right)J. \tag{27}$$

While Eq.(4) or equivalently Eq.(20) tracks the true accumulated volume variation, and the adaptive map results in the modification of volume variation by $\overline{\beta}[\mathrm{tr}(\mathbf{U} - \mathbf{I}) + \mathrm{tr}(\mathbf{F}(\overline{\mathbf{x}} \cdot \overline{\nabla}\mathbf{R}))]J$. This means while the effective accumulated growth is updated by the local growth rate $\overline{\Gamma}$, it is further modified by the relaxation rate $\overline{\beta}$. This modification does not affect the mass conservation, since the mass conservation is enforced by Eq.(3), but it modifies the growth tensor dynamics through Eq.(7). Further, it does not violate the incompressibility condition $J_e = \det(\mathbf{F}_e) = 1$ because of the following. From Eqs. (7), (8), (25) and (27), one can derive the evolution of the elastic deformation gradient

$$\overset{\diamond}{\mathbf{F}}_e = -\left(\overline{\beta}[\mathbf{R}\left(\mathbf{U} - \mathbf{I}\right)\mathbf{R}^T - \frac{1}{3}\mathrm{tr}(\mathbf{U} - \mathbf{I})\mathbf{I}] + \overline{\beta}[\mathbf{F}(\overline{\mathbf{x}} \cdot \overline{\nabla}\mathbf{R}) - \frac{1}{3}\mathrm{tr}(\mathbf{F}(\overline{\mathbf{x}} \cdot \overline{\nabla}\mathbf{R}))] + \frac{\overline{\Gamma}}{3}\mathbf{I}\right)\mathbf{F}_e. \tag{28}$$

One can check $\frac{DJ_e}{Dt} = 0$ from Eq. (28) and thus $J_e = 1$ is maintained.

Notice that Eqs. (27) and (28) are both objective under rotation. Interestingly, only the growth rate $\overline{\Gamma}$ contributes to the diagonal components of the tensor before $\mathbf{F}_e$ on the right hand side of Eq. (28) since both $\overline{\beta}[\mathbf{R}\left(\mathbf{U} - \mathbf{I}\right)\mathbf{R}^T - \frac{1}{3}\mathrm{tr}(\mathbf{U} - \mathbf{I})\mathbf{I}]$ and $\overline{\beta}[\mathbf{F}(\overline{\mathbf{x}} \cdot \overline{\nabla}\mathbf{R}) - \frac{1}{3}\mathrm{tr}(\mathbf{F}(\overline{\mathbf{x}} \cdot \overline{\nabla}\mathbf{R}))]$ are traceless. Finally, we have that the non-pressure part of the elastic stress $\overline{\boldsymbol{\sigma}}_e = \overline{\mu}_1\mathbf{F}_e\mathbf{F}_e^T$ relaxes according to

$$\begin{aligned}\overset{\triangledown}{\overline{\boldsymbol{\sigma}}}_e = -\frac{2}{3}\,\overline{\Gamma}\overline{\boldsymbol{\sigma}}_e - \overline{\beta}[\mathbf{R}\left(\mathbf{U} - \mathbf{I}\right)\mathbf{R}^T - \frac{1}{3}\mathrm{tr}(\mathbf{U} - \mathbf{I})\mathbf{I} + \mathbf{F}(\overline{\mathbf{x}} \cdot \overline{\nabla}\mathbf{R}) - \frac{1}{3}\mathrm{tr}(\mathbf{F}(\overline{\mathbf{x}} \cdot \overline{\nabla}\mathbf{R}))]\overline{\boldsymbol{\sigma}}_e \\ -\overline{\beta}\overline{\boldsymbol{\sigma}}_e[\mathbf{R}\left(\mathbf{U} - \mathbf{I}\right)\mathbf{R}^T - \frac{1}{3}\mathrm{tr}(\mathbf{U} - \mathbf{I})\mathbf{I} + (\overline{\mathbf{x}} \cdot \overline{\nabla}\mathbf{R})^T\mathbf{F}^T - \frac{1}{3}\mathrm{tr}(\mathbf{F}(\overline{\mathbf{x}} \cdot \overline{\nabla}\mathbf{R}))].\end{aligned} \tag{29}$$

Notice that Eq.(29) is also objective under rotation.

This relaxation dynamics is able to provide a steady-state stress distribution when the tumor spheroid reaches an equilibrium size.

In the linear elasticity regime, where the displacement $\overline{\mathbf{u}}$ is small, we have $\mathbf{F} = \left(\overline{\nabla}\overline{\mathbf{Y}}\right)^{-1} \sim \mathbf{I} + \overline{\nabla}\overline{\mathbf{u}}$ and the Finger deformation tensor $\mathbf{C} = \mathbf{FF}^T \sim \mathbf{I} + \overline{\nabla}\overline{\mathbf{u}} + \overline{\nabla}\overline{\mathbf{u}}^T$, where we have dropped terms that are quadratic and higher powers in the displacement. Without growth, we can show that Eqs. (26) and (29) are similar to the upper convected Maxwell (UCM) viscoelastic model (e.g., see [15]) in the leading order, where the Finger deformation tensor $\mathbf{C} = \mathbf{FF}^T$ follows the dynamics $\overset{\triangledown}{\mathbf{C}} = \overline{\beta}_M\left(\mathbf{I} - \mathbf{C}\right)$ and the Cauchy stress tensor $\tilde{\overline{\boldsymbol{\sigma}}}_e$ follows $\beta_M^{-1}\overset{\triangledown}{\tilde{\overline{\boldsymbol{\sigma}}}}_e + \tilde{\overline{\boldsymbol{\sigma}}}_e = 2\overline{\mu}_1\mathbf{D}$ where

$\mathbf{D} = \left(\nabla\mathbf{v} + \nabla\mathbf{v}^T\right)/2$ is the rate of deformation tensor. First, using $\mathbf{U} \sim \mathbf{I} + \frac{1}{2}(\overline{\nabla}\overline{\mathbf{u}} + \overline{\nabla}\overline{\mathbf{u}}^T)$ and $\mathbf{R} \sim \mathbf{I} + \frac{1}{2}(\overline{\nabla}\overline{\mathbf{u}} - \overline{\nabla}\overline{\mathbf{u}}^T)$ we can derive the leading-order relations

$$\begin{aligned}
&\overline{\beta}\left(\mathbf{R}\left(\mathbf{I}-\mathbf{U}\right)\mathbf{R}^T\mathbf{C} + \mathbf{C}\mathbf{R}\left(\mathbf{I}-\mathbf{U}\right)\mathbf{R}^T\right) - \overline{\beta}\left(\mathbf{F}\left(\overline{\mathbf{x}}\cdot\overline{\nabla}\mathbf{R}\right)\mathbf{C} + \mathbf{C}\left(\overline{\mathbf{x}}\cdot\overline{\nabla}\mathbf{R}\right)^T\mathbf{F}^T\right)\\
&\sim -\overline{\beta}\left(\overline{\nabla}\overline{\mathbf{u}} + \overline{\nabla}\overline{\mathbf{u}}^T\right) - \frac{1}{2}\overline{\beta}\left(\left(\overline{\mathbf{x}}\cdot\overline{\nabla}(\overline{\nabla}\overline{\mathbf{u}} - \overline{\nabla}\overline{\mathbf{u}}^T)\right) + \left(\overline{\mathbf{x}}\cdot\overline{\nabla}(\overline{\nabla}\overline{\mathbf{u}} - \overline{\nabla}\overline{\mathbf{u}}^T)\right)^T\right)\\
&\sim \overline{\beta}\left(\mathbf{I}-\mathbf{C}\right) - \frac{1}{2}\overline{\beta}\left(\left(\overline{\mathbf{x}}\cdot\overline{\nabla}(\overline{\nabla}\overline{\mathbf{u}} - \overline{\nabla}\overline{\mathbf{u}}^T)\right) + \left(\overline{\mathbf{x}}\cdot\overline{\nabla}(\overline{\nabla}\overline{\mathbf{u}} - \overline{\nabla}\overline{\mathbf{u}}^T)\right)^T\right)
\end{aligned} \tag{30}$$

from Eq. (26), which reduces to the Finger deformation tensor dynamics in the UCM model when the gradient of the incremental rotations $\frac{1}{2}(\overline{\nabla}\overline{\mathbf{u}} - \overline{\nabla}\overline{\mathbf{u}}^T)$ is small. Similarly, we can derive from Eq.(29)

$$\begin{aligned}
&-\overline{\beta}[\mathbf{R}\left(\mathbf{U}-\mathbf{I}\right)\mathbf{R}^T - \frac{1}{3}\mathrm{tr}(\mathbf{U}-\mathbf{I})\mathbf{I} + \mathbf{F}(\overline{\mathbf{x}}\cdot\overline{\nabla}\mathbf{R}) - \frac{1}{3}\mathrm{tr}(\mathbf{F}(\overline{\mathbf{x}}\cdot\overline{\nabla}\mathbf{R}))]\overline{\boldsymbol{\sigma}}_e\\
&-\overline{\beta}\overline{\boldsymbol{\sigma}}_e[\mathbf{R}\left(\mathbf{U}-\mathbf{I}\right)\mathbf{R}^T - \frac{1}{3}\mathrm{tr}(\mathbf{U}-\mathbf{I})\mathbf{I} + (\overline{\mathbf{x}}\cdot\overline{\nabla}\mathbf{R})^T\mathbf{F}^T - \frac{1}{3}\mathrm{tr}(\mathbf{F}(\overline{\mathbf{x}}\cdot\overline{\nabla}\mathbf{R}))]\\
&\sim -\overline{\beta}\left(\overline{\nabla}\overline{\mathbf{u}} + \overline{\nabla}\overline{\mathbf{u}}^T\right) - \frac{1}{2}\overline{\beta}\left(\left(\overline{\mathbf{x}}\cdot\overline{\nabla}(\overline{\nabla}\overline{\mathbf{u}} - \overline{\nabla}\overline{\mathbf{u}}^T)\right) + \left(\overline{\mathbf{x}}\cdot\overline{\nabla}(\overline{\nabla}\overline{\mathbf{u}} - \overline{\nabla}\overline{\mathbf{u}}^T)\right)^T\right) + \frac{2}{3}\overline{\beta}\mathrm{tr}\left(\overline{\mathbf{x}}\cdot\overline{\nabla}(\overline{\nabla}\overline{\mathbf{u}} - \overline{\nabla}\overline{\mathbf{u}}^T)\right)^T)\\
&\sim \overline{\beta}\left(\overline{\mu}_1\mathbf{I} - \overline{\boldsymbol{\sigma}}_e\right) - \frac{1}{2}\overline{\beta}\left(\left(\overline{\mathbf{x}}\cdot\overline{\nabla}(\overline{\nabla}\overline{\mathbf{u}} - \overline{\nabla}\overline{\mathbf{u}}^T)\right) + \left(\overline{\mathbf{x}}\cdot\overline{\nabla}(\overline{\nabla}\overline{\mathbf{u}} - \overline{\nabla}\overline{\mathbf{u}}^T)\right)^T\right) + \frac{2}{3}\overline{\beta}\mathrm{tr}\left(\overline{\mathbf{x}}\cdot\overline{\nabla}(\overline{\nabla}\overline{\mathbf{u}} - \overline{\nabla}\overline{\mathbf{u}}^T)\right)^T),
\end{aligned} \tag{31}$$

where we have used incompressibility $\det(\mathbf{F}) = 1$ ($\overline{\nabla}\cdot\overline{\mathbf{u}} = 0$). Assuming the gradient of the incremental rotations are negligible, the non-pressure part of the stress tensor $\overline{\boldsymbol{\sigma}}_e = \overline{\mu}_1\mathbf{C}$ relaxes according to $\beta_M^{-1}\overset{\triangledown}{\overline{\boldsymbol{\sigma}}}_e + \overline{\boldsymbol{\sigma}}_e = \overline{\mu}_1\mathbf{I}$ at the leading order. We can define the Cauchy stress tensor $\tilde{\overline{\boldsymbol{\sigma}}}_e := \beta_M\left(\overline{\boldsymbol{\sigma}}_e - \overline{\mu}_1\mathbf{I}\right) = \overline{\mu}_1\beta_M\left(\mathbf{C}-\mathbf{I}\right)$ and show $\beta_M^{-1}\overset{\triangledown}{\tilde{\overline{\boldsymbol{\sigma}}}}_e + \tilde{\overline{\boldsymbol{\sigma}}}_e = 2\overline{\mu}_1\mathbf{D}$ (where we have used $\overset{\triangledown}{\mathbf{I}} = -2\mathbf{D}$), which is the UCM model of viscoelasticity. We note that in the context of tumor growth, there are other models of stress relaxation that have been used that also reduce to Maxwell viscoelastic models in the limit of small displacements, e.g., see [16, 17].

The relaxation proposed in Eq. (23) is similar in spirit to the approach used in [18] to model viscoelasticity in the context of a porous viscoelastic model for the cell cytoskeleton where the reference frame also relaxes to the current configuration but instead [18] uses the Lagrangian frame and a different formulation. In [18], the reference configuration is written as $\overline{\mathbf{s}}(\overline{\boldsymbol{\alpha}},\overline{t}) = \overline{\mathbf{Y}}(\overline{\mathbf{x}}(\overline{\boldsymbol{\alpha}},\overline{t}),\overline{t})$ where $\overline{\boldsymbol{\alpha}}$ denotes a particle label and $\frac{\partial}{\partial\overline{t}}\overline{\mathbf{x}}(\overline{\boldsymbol{\alpha}},\overline{t}) = \overline{\mathbf{v}}(\overline{\mathbf{x}}(\overline{\boldsymbol{\alpha}},\overline{t}),\overline{t})$ is the velocity of the viscoelastic structure in the current frame. Instead of using Eq. (23), the relaxation in [18] is implemented as $\mathbf{F}\frac{\partial\overline{\mathbf{s}}}{\partial\overline{t}} = \overline{\beta}_G\left(\overline{\mathbf{x}}(\overline{\boldsymbol{\alpha}},\overline{t}) - \overline{\mathbf{s}}(\overline{\boldsymbol{\alpha}},\overline{t})\right)$, where $\overline{\beta}_G$ is the relaxation rate. This is designed to model the relaxation $\overset{\diamond}{\mathbf{F}} = \overline{\beta}_G\left(\mathbf{I}-\mathbf{F}\right)$ approximately by dropping terms involving spatial derivatives of $\mathbf{F}$ other than the advection term $\overline{\mathbf{v}}\cdot\overline{\nabla}\mathbf{F}$, see [18] for details.

#### *1.1.5 Initial, boundary conditions and the moving boundary*

Consider a tissue region $\overline{\Omega}_{\overline{t}}$ that expands with a moving boundary $\overline{\Sigma}_{\overline{t}}$. Assuming the tissue is initially stress-free, the initial condition for $\overline{\mathbf{Y}}$ from Eq.(23) at $\overline{t} = \overline{0}$ is

$$\overline{\mathbf{Y}}(\overline{\mathbf{x}},\overline{0}) = \overline{\mathbf{x}}, \overline{\mathbf{x}} \in \overline{\Omega}_0. \tag{32}$$

The traction balance at the moving boundary gives

$$\overline{\boldsymbol{\sigma}}\hat{\mathbf{n}}|_{\overline{\Sigma}_{\overline{t}}} = \overline{\mathbf{T}} \tag{33}$$

$$\tag{34}$$

where $\hat{\mathbf{n}}$ is the outward unit normal and $\overline{\mathbf{T}}$ is the traction at the boundary. When $\overline{\mathbf{T}} = \overline{\mathbf{0}}$ there is no external compression (or surface tension). The dynamics of the boundary is given by

$$\overline{V} = \overline{\mathbf{v}}\cdot\hat{\mathbf{n}}|_{\overline{\Sigma}_{\overline{t}}} \tag{35}$$

where $\overline{\mathbf{v}}$ is the velocity field from Eq. (1).

#### 1.2. The non-dimensionalized system

Here we non-dimensionalize the equations (1),(3),(11), and (23) by taking $\overline{\mathbf{x}} \to \mathbf{x}l$, $\overline{t} \to t\tau$, $\overline{\mathbf{v}} \to \mathbf{v}l/\tau$, $\overline{\tilde{\boldsymbol{\sigma}}} \to \tilde{\boldsymbol{\sigma}}\overline{\mu}_1$, $\overline{p} \to p\overline{\mu}_1$, $\overline{\Gamma}(\overline{\mathbf{x}}, \overline{t}) \to \Gamma(\mathbf{x}, t)/\tau$, $\overline{\beta} \to \beta/\tau$ and $\overline{\alpha} \to \alpha\overline{\mu}_1\tau/l^2$, where $l$ is a characteristic length, $\tau$ is a characteristic time and the unbarred quantities are dimensionless. Here, we take $l = 1\mu$m and $\tau = 1$ day. We further assume that the drag is small relative to the shear stress: $\overline{\alpha}/\overline{\mu}_1 = 0.001$ day/$\mu$m$^2$, which implies that the deviation from elastic equilibrium is small. The nonlinear dimensionless system is the following:

$$\nabla \cdot \mathbf{v} = \Gamma(\mathbf{x}, t) \tag{36}$$

$$\alpha \mathbf{v}(\mathbf{x}, t) = \nabla \cdot \tilde{\boldsymbol{\sigma}} \tag{37}$$

$$\tilde{\boldsymbol{\sigma}}(\mathbf{x}, t) = [\det(\nabla \mathbf{Y})]^{2/3}[\nabla \mathbf{Y}]^{-1}[\nabla \mathbf{Y}]^{-1T} + \mu[\det(\nabla \mathbf{Y})]^{-2/3}[\nabla \mathbf{Y}]^T[\nabla \mathbf{Y}] - p\mathbf{I}. \tag{38}$$

$$\mathbf{Y}_t + \mathbf{v} \cdot \nabla \mathbf{Y} = \beta(\mathbf{x} - \mathbf{Y}) \tag{39}$$

where $\mu = \overline{\mu}_2/\overline{\mu}_1$ and $\mathbf{Y}_t = \frac{\partial \mathbf{Y}}{\partial t}$. The nondimensional initial and boundary conditions become

$$\mathbf{Y}(\mathbf{x}, 0) = \mathbf{x}, \mathbf{x} \in \Omega_0, \tag{40}$$

$$\tilde{\boldsymbol{\sigma}}\hat{\mathbf{n}}|_{\Sigma_t} = \mathbf{T}, \tag{41}$$

$$V = \mathbf{v} \cdot \hat{\mathbf{n}}|_{\Sigma_t}, \tag{42}$$

where $\overline{\Omega}_0 \to \Omega_0 l^3$, $\overline{\Omega}_{\overline{t}} \to \Omega_t l^3$, $\overline{\Sigma}_{\overline{t}} \to \Sigma_t l^2$, $\mathbf{T} = \overline{\mathbf{T}}/\overline{\mu}_1$ and $V = \overline{V}\tau/l$.

To compare with previous results accounting for linear elasticity, here we change $\mathbf{Y} = \mathbf{x} - \mathbf{u}$ where $\mathbf{u}$ is the displacement field. Then Eq.(39) can be rewritten as

$$\mathbf{u}_t + \mathbf{v} \cdot \nabla \mathbf{u} = \mathbf{v} - \beta \mathbf{u}. \tag{43}$$

When $\mathbf{u}$ is small: $|\mathbf{u}| \sim \epsilon$, Eq.(38) can be linearized to

$$\tilde{\boldsymbol{\sigma}}(\mathbf{x}, t) = (1 - \mu)[(\nabla \mathbf{u} + \nabla \mathbf{u}^T) - \frac{2}{3}(\nabla \cdot \mathbf{u})\mathbf{I})] - p\mathbf{I}, \tag{44}$$

which is a linear elastic constitutive equation between stress and strain with pressure as a Lagrange multiplier to ensure incompressibility. The boundary condition (41) can be rewritten as

$$(1 - \mu)[(\nabla \mathbf{u} + \nabla \mathbf{u}^T) - \frac{2}{3}(\nabla \cdot \mathbf{u})\mathbf{I})]\mathbf{n}|_{\Sigma_t} - p\mathbf{n}|_{\Sigma_t} = \mathbf{T}, \tag{45}$$

where we have implicitly assumed that $(\mathbf{I} - \hat{\mathbf{n}} \otimes \hat{\mathbf{n}})\,\mathbf{T} \sim O(\epsilon)$ because if this is not the case then Eq. (45) does not balance unless $1 - \mu \sim 1/\epsilon$.

Linear elastic (or more generally poroelastic/mixture) models have been considered in a number of previous works including Jones et al. [11], Roose et al. [19], Araujo and McElwain [12], Ambrosi and Preziosi [20]. In these references, the (material) time derivative was taken to relate the rate of change of stress to the velocity field and strain rate. In particular, the time derivatives of $\nabla \cdot \mathbf{u}$ and the strain tensor $1/2(\nabla \mathbf{u} + \nabla \mathbf{u}^T)$ become the volumetric growth rate $\nabla \cdot \mathbf{v}$ and rate of strain tensor $1/2(\nabla \mathbf{v} + \nabla \mathbf{v}^T)$, respectively, since $\frac{d\mathbf{u}}{dt} = \mathbf{v}$ where $d/dt = \partial_t + \mathbf{v} \cdot \nabla$ is the material time derivative. When remodeling via Eq. (43) is considered, $\mathbf{u}$ is no longer the traditional displacement field and instead $\frac{d\mathbf{u}}{dt} = \mathbf{v} - \beta \mathbf{u}$. Then combining Eqs. (37), (43) and (44), the linear approximation becomes

$$-\alpha \mathbf{v}(\mathbf{x}, t) + (1 - \mu)\nabla \cdot [(\nabla \mathbf{u} + \nabla \mathbf{u}^T) - \frac{2}{3}(\nabla \cdot \mathbf{u})\mathbf{I})] - \nabla p = 0. \tag{46}$$

However, when $\beta$ is small and there is significant accumulated volumetric growth, and the displacement $\mathbf{u}$ may not be small, which would make the linear approximation inaccurate.

#### *1.2.1 Fluid models*

There are two ways to obtain a fluid model as the leading order limit of our system. In the first, observe that when the boundary traction $|\mathbf{T}\hat{\mathbf{n}}| \sim O(1)$ and $|\mathbf{u}| \sim \epsilon$, the leading order of the boundary condition equation (45) becomes

$$-p|_{\Sigma_t} = \mathbf{T}\hat{\mathbf{n}} \tag{47}$$

which results in $p \sim O(1)$ at leading order since the elastic stresses are $O(\epsilon)$. When $\alpha \sim O(1)$, we conclude that at leading order Eq. (1) reduces to Darcy's model (see Eq. (50) below). When $\alpha \sim O(\epsilon)$, a trivial case of uniform pressure determined by the boundary traction $\mathbf{T}$ is obtained. In both cases, the displacement field $\mathbf{u}$ does not contribute to the leading order stress.

When the boundary traction $|\mathbf{T}\hat{\mathbf{n}}| \sim O(\epsilon)$, we can still obtain a fluid model when the growth rate $\Gamma(\mathbf{x}, t) \sim O(1)$ is finite and the remodeling rate $\beta \sim O(1/\epsilon)$ is large. It follows from Eq. (36) that $|\mathbf{v}(\mathbf{x}, t)| \sim O(1)$ and from Eq. (43) that $|\mathbf{u}(\mathbf{x}, t)| \sim O(\epsilon)$ and

$$\mathbf{v} = \beta\mathbf{u} \tag{48}$$

at leading order. This makes Eq.(46) become

$$-\alpha\mathbf{v}(\mathbf{x}, t) + \frac{(1-\mu)}{\beta}\nabla \cdot [(\nabla\mathbf{v} + \nabla\mathbf{v}^T) - \frac{2}{3}(\nabla \cdot \mathbf{v})\mathbf{I})] - \nabla p = 0, \tag{49}$$

where $\frac{(1-\mu)}{\beta} \sim \epsilon$ is the effective viscosity. At leading order, we recover

1. Darcy's model
$$-\alpha\mathbf{v}(\mathbf{x}, t) - \nabla p = 0 \tag{50}$$
when $\alpha \sim O(1)$;
2. Darcy-Stokes model, e.g. a rescaled version of Eq.(49), when $\alpha \sim O(\epsilon)$;
3. Stokes' model
$$\frac{(1-\mu)}{\beta}\nabla \cdot [(\nabla\mathbf{v} + \nabla\mathbf{v}^T) - \frac{2}{3}(\nabla \cdot \mathbf{v})\mathbf{I})] - \nabla p = 0, \tag{51}$$
when $\alpha \sim O(\epsilon^2)$.

In all the linearized cases, we can solve $\mathbf{v}$ and $p$ first from Eqs. (36) and (49). Then, $\mathbf{u}$ can be computed from Eq. (48).

#### *1.2.2 Linear elastic models*

When the boundary traction $|\mathbf{T}| \sim O(\epsilon)$, the growth rate $\Gamma(\mathbf{x}, t) \sim O(\epsilon)$, the relaxation rate $\beta \sim O(1)$, it follows that $|\mathbf{v}(\mathbf{x}, t)| \sim O(\epsilon)$ and $|\mathbf{u}(\mathbf{x}, t)| \sim O(\epsilon)$ and

$$\mathbf{u}_t = \mathbf{v} - \beta\mathbf{u} \tag{52}$$

at the leading order. Then, we can rewrite Eq.(36) as

$$(\nabla \cdot \mathbf{u})_t = \Gamma(\mathbf{x}, t) - \beta(\nabla \cdot \mathbf{u}) \tag{53}$$

and Eq.(46) as

$$-\alpha(\mathbf{u}_t + \beta\mathbf{u}) + (1-\mu)\nabla \cdot [(\nabla\mathbf{u} + \nabla\mathbf{u}^T) - \frac{2}{3}(\nabla \cdot \mathbf{u})\mathbf{I})] - \nabla p = 0, \tag{54}$$

where the linear elastic stress balances the pressure and drag. Thus, at leading order, we obtain

1. A linear elastic model with drag Eq.(54) when $\alpha \sim O(1)$;

2. A linear elastic model

$$(1-\mu)\nabla\cdot[(\nabla\mathbf{u}+\nabla\mathbf{u}^T)-\frac{2}{3}(\nabla\cdot\mathbf{u})\mathbf{I})]-\nabla p=0 \tag{55}$$

when $\alpha\sim O(\epsilon)$.

In both cases, we solve $\mathbf{u}$ and $p$ together from Eqs. (53) and (54) and afterward, $\mathbf{v}$ can be solved from Eq.(52).

#### 1.3. The radially symmetric case: Growth of tumor spheroids

In radial coordinates, Eqs. (36)-(39) reduce to

$$v(r,t)=\frac{1}{r^2}\int_0^r\Gamma(\eta,t)\eta^2 d\eta \tag{56}$$

$$p(r,t)_{,r}=-\alpha v(r,t)+\sigma_{rr}(r,t)_{,r}+\frac{2}{r}(\sigma_{rr}(r,t)-\sigma_{\theta\theta}(r,t)) \tag{57}$$

$$\sigma_{rr}(r,t)=(\frac{y/r}{y_{,r}})^{4/3}+\mu(\frac{y/r}{y_{,r}})^{-4/3}\text{ and }\sigma_{\theta\theta}(r,t)=(\frac{y_{,r}}{y/r})^{2/3}+\mu(\frac{y_{,r}}{y/r})^{-2/3} \tag{58}$$

$$y(r,t)_{,t}+v(r,t)y(r,t)_{,r}=\beta(r-y(r,t)). \tag{59}$$

with the initial condition

$$y(r,0)=r \tag{60}$$

and the boundary condition

$$p(R,t)-(\frac{\frac{y}{r}}{y_{,r}})^{4/3}|_{r=R}-\mu(\frac{\frac{y}{r}}{y_{,r}})^{-4/3}|_{r=R}=F_{compression}. \tag{61}$$

The radius evolves via

$$\frac{dR}{dt}=v(R,t) \tag{62}$$

given $R(0)=R_0$. When the tumor is embedded in a gel, $F_{compression}$ can be calculated analytically assuming there is no drag in the gel and that the gel does not remodel (see Eq.(76) below). To solve the system numerically, we can convert the above moving-boundary problem to a fixed-domain problem. Introducing the rescaled coordinate $r'$ such that $r\to R(t)r'$, the derivatives with respect to $t$ and $r$ above can be rewritten as

$$\Box_{,t}=\tilde{\Box}_{,t}-r'\frac{\dot{R}}{R}\tilde{\Box}_{,r'} \tag{63}$$

$$\Box_{,r}=\frac{1}{R}\tilde{\Box}_{,r'}. \tag{64}$$

and we solve the converted system in the domain $0\le r'\le 1$, which corresponds to $0\le r\le R(t)$ at each time step.

We assume that the nondimensional growth rate is given by

$$\Gamma(r,t)=\lambda(r,t)-\lambda_A(r,t) \tag{65}$$

where $\lambda(r,t)$ is the volume gain (e.g., due to cell proliferation) and $\lambda_A(r,t)$ is the volume loss (e.g., due to cell death and cell compaction). We assume that cells proliferate in response to the presence of nutrients, which we model as [11, 21, 22]

$$\lambda(r,t)=\lambda_0 c(r,t) \tag{66}$$

where $\lambda_0$ is a rate and $c(r,t)$ is the concentration of nutrients. Following [11, 21, 22], we assume that $c$ satisfies

$$L^2\Delta c-c=0\text{ in }\Omega_t, \tag{67}$$

$$c=1\text{ on }\Sigma_t. \tag{68}$$

In radial coordinates the solution is given by

$$c(r,t) = (R(t)\sinh(\frac{r}{L}))/(r\sinh(\frac{R(t)}{L})). \tag{69}$$

We assume that compressive stress induces volume loss due to increased cell death and compression of cells. Accordingly, we take

$$\lambda_A(r,t) = \lambda_{A,0} + \Delta_A \frac{\gamma_A(\sigma_{\theta\theta}-p)^n \cdot 1_{\{\sigma_{\theta\theta}-p<0\}}}{1+\gamma_A(\sigma_{\theta\theta}-p)^n}. \tag{70}$$

At each time step $n$, we solve the nonlinear system as follows:

1. Step 1: Given $y^n$, update $\sigma_{rr}^n$ and $\sigma_{\theta\theta}^n$ from Eq. (58).

2. Step 2: Given $\sigma_{rr}^n$ and $\sigma_{\theta\theta}^n$, solve the coupled equations for the velocity $v^n$ and pressure $p^n$: $v^n = \frac{1}{r^2}\int_0^r (\Gamma(\sigma_{\theta\theta}^n, p^n)\eta^2 d\eta$ and $p_{,r}^n = -\alpha v^n + \sigma_{rr,r}^n + \frac{2}{r}(\sigma_{rr}^n - \sigma_{\theta\theta}^n)$, by first discretizing the equations and then solving the nonlinear discrete system using the nonlinear solver "fsolve" in Matlab .

3. Step 3: Given $y^n$ and $v^n$, Solve $y^{n+1}$ from $(y^{n+1}-y^n)/\Delta t + v^n y_{,r}^n = \beta(r - y^{n+1})$ where $y_{,r}^n$ uses the 1st order upwind scheme.

The numerical scheme has been validated using convergence tests and through comparison with the analytical solution provided below in Sec. 3.2.

##### *1.3.1 Radial compression from the surrounding gel*

The traction at the boundary of the spheroid can arise from surface tension and externally applied stress. For example, when the spheroid is in a fluid, $F_{compression} = p_{ext}$ where $p_{ext} = -\gamma_{st}\kappa + p_{fluid}$ where $\gamma_{st}$ is the surface tension coefficient and $p_{fluid}$ is the pressure in the fluid, e.g. generated by osmotic stress [23]. If $p_{ext} = 0$, then there is no applied traction. When the spheroid is placed into a gel, the gel can exert stress on the spheroid. Suppose that the exterior gel is incompressible, isotropic and neo-Hookean (e.g., $\overline{\mu}_2 = 0$ in Eq. (11)) and nondimensional shear modulus $c_H$ (ratio of actual shear modulus of the gel and the tumor spheroid). We also assume there is no drag in the gel ($\alpha = 0$), there is no production of new extracellular material ($\Gamma = 0$) and that the gel does not remodel ($\beta = 0$). Then, solving the nondimensional system (56)-(62) replacing the boundary condition at $r = R$ with a traction boundary condition at $r = \infty$, we obtain

$$y(r,t) = (r^3 - R(t)^3 + R_0^3)^{\frac{1}{3}} \tag{71}$$

$$v(r,t) = \frac{R^2}{r^2}\frac{dR}{dt} \tag{72}$$

$$\sigma_{ext,rr} = c_H \frac{(r^3 - R(t)^3 + R_0^3)^{\frac{4}{3}}}{r^4} \tag{73}$$

$$\sigma_{ext,\theta\theta} = c_H \frac{r^2}{(r^3 - R(t)^3 + R_0^3)^{\frac{2}{3}}} \tag{74}$$

$$p_{ext}(r,t) = c_H \frac{(r^3 - R^3 + R_0^3)^{\frac{4}{3}}}{r^4} - c_H \frac{(r^3 - R^3 + R_0^3)^{\frac{1}{3}}(5r^3 + R_0^3 - R^3)}{2r^4} + c_p \tag{75}$$

where $c_p$ is a constant determined by the far field ($r = \infty$) boundary condition for the stress. For example, if there is no applied traction at $r = \infty$, then $c_p = (5/2)c_H$. Assuming this is the case, the compression from the gel at the moving boundary is:

$$F_{compression} = -(\sigma_{ext,rr} - p_{ext})|_{r=R} = \frac{c_H}{2}\left(5 - \frac{R_0(R_0^3 + 4R^3)}{R^4}\right). \tag{76}$$

*1.3.2 Analytical solutions when $\Gamma(r,t) = const.$*

When $\Gamma(r,t) = \Gamma$ is a constant, the analytical solution to Eqs. (56)-(62) is

$$v(r,t) = \frac{\Gamma r}{3} \tag{77}$$

$$y(r,t) = \frac{r}{\beta + \frac{\Gamma}{3}}\left(\beta + \frac{\Gamma}{3} e^{-(\beta + \frac{\Gamma}{3})t}\right) \tag{78}$$

$$p(r,t) = 1 + \mu + \frac{\alpha\Gamma}{6}(R^2 - r^2) + F_{compression}. \tag{79}$$

$$\tag{80}$$

Notice the elastic stresses, $\sigma_{rr}(r,t) = \sigma_{\theta\theta}(r,t) = 1 + \mu$, are constant. Although this case is not biologically meaningful (e.g., uniform growth), this case is useful for helping to validate the numerical method.

### 1.4 Supplemental figures

### 1.5 Glossary

| | |
|---|---|
| $\sigma_{rr}$ | non-dimensional radial elastic stress |
| $\sigma_{\theta\theta}$ | non-dimensional circumferential/hoop elastic stress |
| $\sigma_{rr}^{tot}$ | total radial elastic stress |
| $\sigma_{\theta\theta}^{tot}$ | total circumferential/hoop elastic stress |
| $\mu$ | rescaled shear modulus |
| $p$ | non-dimensional pressure |
| $R$ | tumor radius |
| $F_{ext}$ | non-dimensional external compression |
| $y$ | initial radial position of a material point |
| $v$ | radial velocity |
| $\lambda$ | the rate of volume increase |
| $\lambda_A$ | the rate of volume loss |
| $\lambda_E$ | the rate of water efflux |
| $\lambda_{net}$ | local volumetric growth rate |
| $c$ | nutrient level |
| $\lambda_c$ | rate of cell divisions |
| $\lambda_{A,c}$ | rate of apoptosis |
| $\rho_c$ | density of cell numbers |

**Parameters for Figure 1**

| | |
|---|---|
| $R_0 = 75$ | tumor initial radius (in $\mu$m) |
| $L = 40$ | diffusional length of nutrient (in $\mu$m) |
| $\beta = 0.6$ | relaxation rate |
| $c_H = 0$ | nondimensional shear modulus |
| $\lambda_0 = 1.1$ | rate of cell volume growth |
| $\lambda_{A,c} = 0.2$ | rate of apoptosis |
| $\Delta_A = 0.9$ | maximum rate of the water flux, |
| $\gamma_A = 0.1$ | sensitivity to the feedback on $\lambda_E$ |
| $\gamma_c = 0.1$ | sensitivity to the feedback on $\lambda_c$ |
| $n = 2$ | Hill coefficient of feedback on $\lambda_E$ |
| $l = 2$ | Hill coefficient of feedback on $\lambda_c$ |

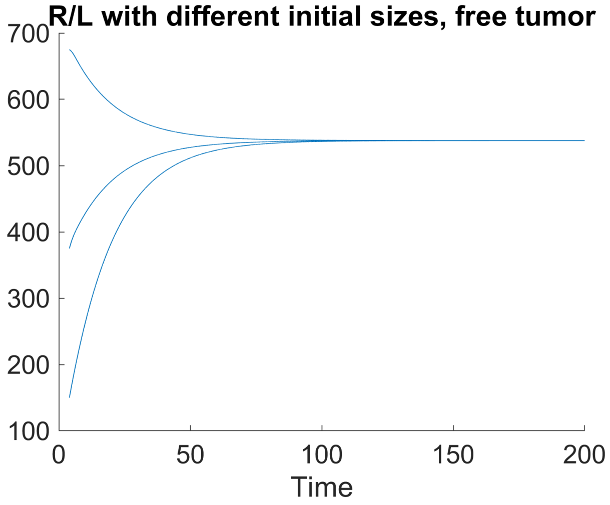


Fig.S 2: Time evolution of the tumor radius with different initial size ($c_H = 0$).

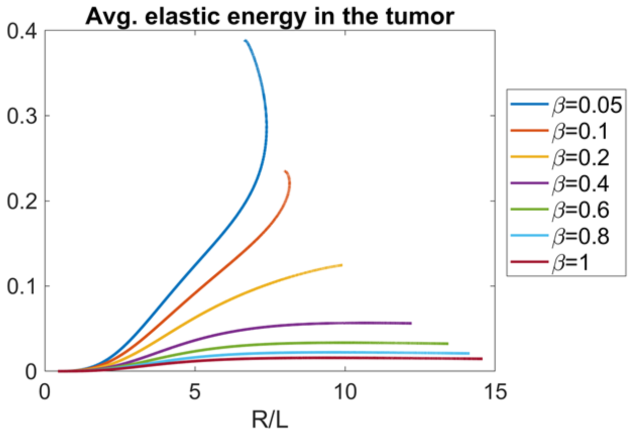


Fig.S 3: Time evolution of the average elastic energy in the tumor, $\frac{3}{4R^3}\int_0^R W(r)r^2 dr$, for different $\beta$. Larger $\beta$ changes the shape of average density and decreases the magnitude due to stress relaxation ($c_H = 0$).

[1] Y. Fung, *Motion, Flow, Stress and Growth* (Springer-Verlag, 1990).
[2] R. W. Ogden, *Non-linear elastic deformations* (Courier Corporation, 1997).
[3] G. A. Holzapfel, Meccanica **37**, 489 (2002).
[4] E. K. Rodriguez, A. Hoger, and A. D. McCulloch, Journal of biomechanics **27**, 455 (1994).
[5] H.-G. Cottet, E. Maitre, and T. Milcent, ESAIM **42**, 471 (2008).
[6] K. Kamrin, C. Rycroft, and J. Nave, J. Mech. Phys. Sol. **60**, 1952 (2012).
[7] M. B. Amar and A. Goriely, Journal of the Mechanics and Physics of Solids **53**, 2284 (2005).
[8] A. Goriely and M. B. Amar, Physical review letters **94**, 198103 (2005).
[9] A. Goriely, *The mathematics and mechanics of biological growth*, vol. 45 (Springer, 2017).
[10] D. Ambrosi and F. Mollica, International journal of engineering science **40**, 1297 (2002).
[11] A. Jones, H. Byrne, J. Gibson, and J. Dold, Journal of mathematical biology **40**, 473 (2000).
[12] R. P. Araujo and D. S. McElwain, SIAM Journal on Applied Mathematics **66**, 447 (2005).
[13] S. Lubkin and T. Jackson, Journal of biomechanical engineering **124**, 237 (2002).
[14] G. Haller, Journal of the Mechanics and Physics of Solids **86**, 70 (2016).
[15] D. Joseph, *Fluid dynamics of viscoelastic fluids* (Springer-Verlag, 1990).

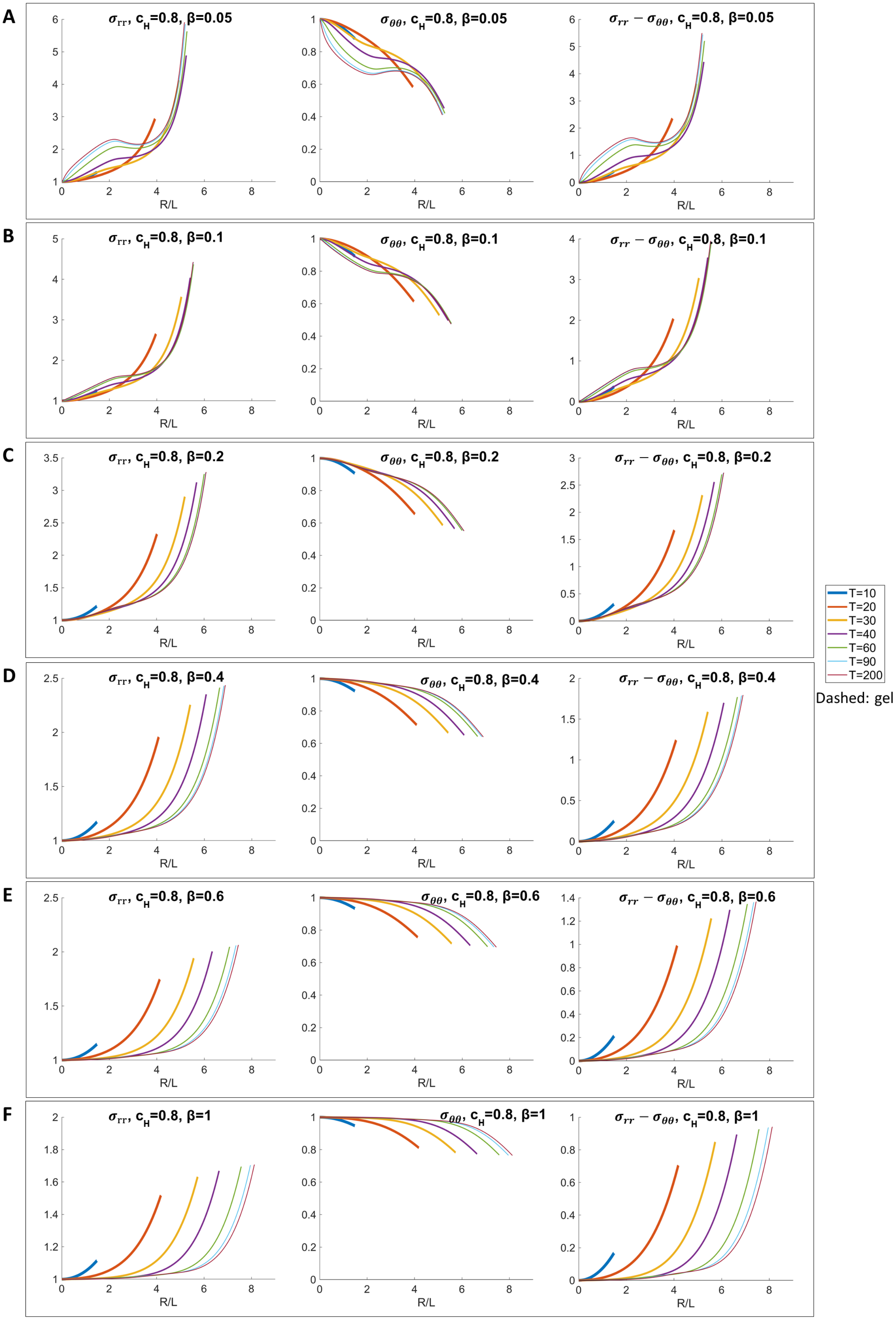


Fig.S 4: Plots of the radial stress ($\sigma_{rr}$), circumferential stress ($\sigma_{\theta\theta}$), and stress anisotropy ($\sigma_{rr} - \sigma_{\theta\theta}$) with $c_H = 0.8$ and different $\beta$. Larger $\beta$ changes the shape of stress, reduces the magnitude and results in more homogeneous stress.

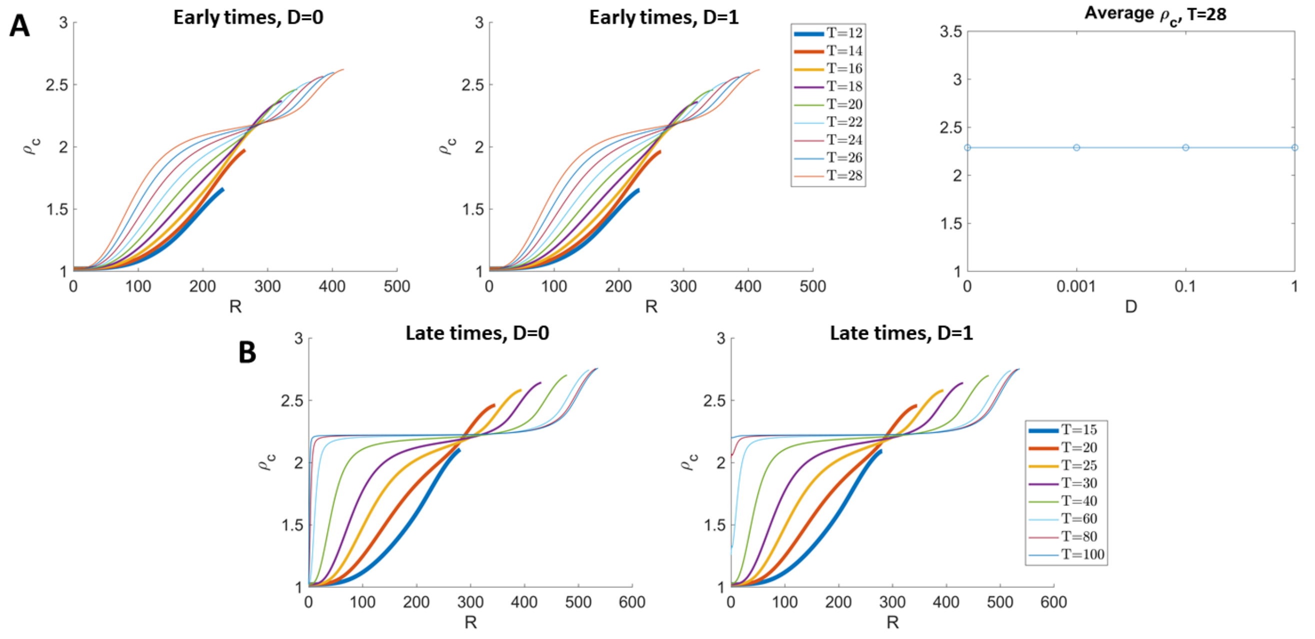


Fig.S 5: Effect of the cell number flux due to local cell neighbor exchanges, modeled by parameter $D$ in Eq. 6 in the main text. Changing $D$ does not affect the distribution $\rho_c$ by T=28, which is the time point of data in [24]. Average $\rho_c$ in the tumor is not affected by $D$ either. At later times, $\rho_c(0,t)$ is a constant due to our numerical scheme unless $D > 0$ (panel B).
