## Supplemental Data for Figure 2 for "Stress generation, relaxation and size control in confined tumor growth"

(Dated: August 30, 2021)

### 2. SUPPLEMENTAL DATA FOR FIGURE 2

In [1], it was shown that tumor spheroids generated with the same cell lines reach different equilibrium sizes when suspended in gels with different stiffness generated by different gel concentrations. Further, if the gels are removed enzymatically, the tumors resume growing to their free suspension sizes. Here, we fit the growth data of tumors in free suspension and in 0.3%, 0.7%, 0.9% and 1.0% agarose gels in Fig. 1A in [1] using our mathematical model. We only consider the feedback on cell death (and compression), as suggested in [1]. We assume that $\lambda$, $\lambda_{A,0}$, $L$, $\beta$, $\Delta_A$, and $\gamma_A$ are identical for a given cell line and only $c_H$ is allowed to vary when the stiffness of the gel is changed. The shear modulus $c_H = 0$ when the tumor is in free suspension (termed "free"). We first fit the free, 0.7% and 1.0% tumors together, then fit $c_H$ for other tumors.

We perform a grid search of parameters to minimize the $l^2$-error of the model using data extracted from [1]. The parameter combination that minimizes the sum of the $l^2$-error of the model is:

$$\begin{aligned}
&\lambda_0 = 1.1, L = 40, \lambda_{A,0} = 0.2, \Delta_A = 0.9, \gamma_A = 0.1, n = 2, \beta = 0.6 \\
&\text{Free: } c_H = 0 \\
&0.3\%\text{: } c_H = 1.0 \\
&0.7\%\text{: } c_H = 1.2 \\
&0.9\%\text{: } c_H = 2.3 \\
&1.0\%\text{: } c_H = 3.4
\end{aligned}$$

The best fits (curves) are shown in Figure S1A. The 0.3%, 0.7%, and 0.9% tumors are in good agreement with the experimental data (symbols). The shaded regions correspond to results using parameters within 10% of the best fit (see Figure S1A and the description below). Increasing the stiffness of the gel makes the tumor more compressive, and reduces the magnitude of circumferential stress outside the tumor (Figure S1B-G). Accordingly, the elastic energy density

$$W(r, y, y_{,r}) = \frac{1}{2}\left(\left(\frac{y/r}{y_{,r}}\right)^{4/3} + 2\left(\frac{y_{,r}}{y/r}\right)^{2/3} - 3\right), \tag{1}$$

both inside and outside the tumor spheroid, and the average elastic energy

$$\frac{3}{4R_\infty^3}\int_0^{R_\infty} W(r, y, y_{,r}) r^2 dr, \tag{2}$$

are also reduced by the gel stiffness (Figure S2).

Next, we fit the data without feedback on $\lambda_A$ from mechanical stresses. The tumor radius $R(t)$ is determined by $\lambda$, $\lambda_{A_0}$ and $L$. In this case, the recovery experiments cannot be reproduced without simultaneously changing all the parameters. Nevertheless, experiments in Figure 2A can be fit by the parameter combinations that minimize the $l^2$-error below:

| | $\lambda_0$ | $\lambda_A$ | $L$ |
|---|---|---|---|
| Free | 0.95 | 0.3 | 60 |
| 0.7% | 1.0 | 0.35 | 30 |
| 1.0% | 1.0 | 0.5 | 8 |

Observe that $\lambda_0$ is nearly constant across the three configurations but $\lambda_A$ is an increasing function of gel concentration and $L$ is a decreasing function of gel concentration.

We use the corrected Akaike information criterion (AICc) [2] to determine the model that better fits the data. The model with a lower AICc score provides better explanation for the data. The AICc score is defined as

$$AICc = n\log\left(\frac{RSS}{n}\right) + \frac{2Kn}{n-K-1},$$

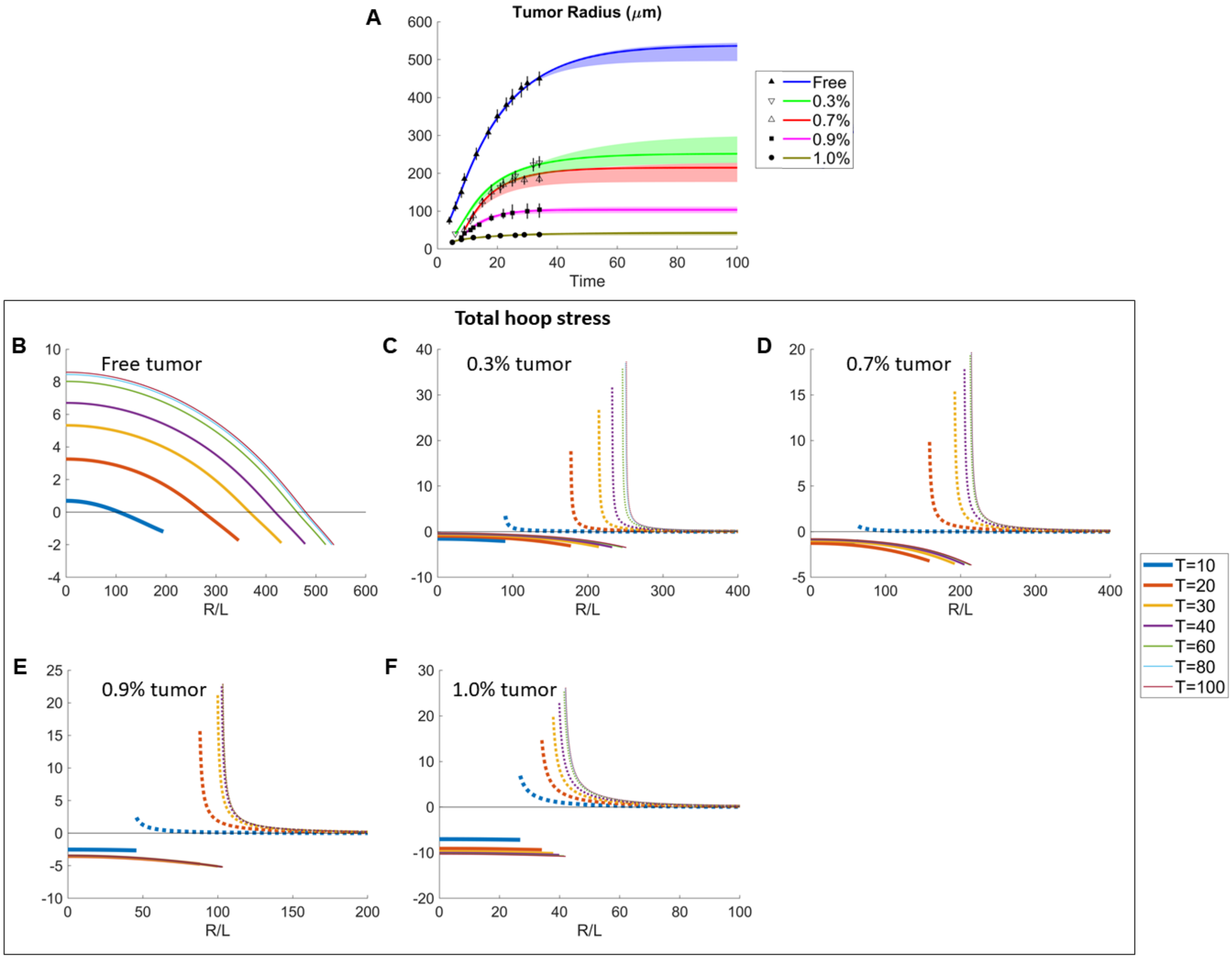


Fig.S 1: (A) Fitting data of the tumor size evolution in free suspension, and in 0.3%, 0.7%, 0.9% and 1.0% agarose gels from Fig. 1A in [1]. (B-G) Time evolution of the total circumferential stress inside the tumor (solid lines) and outside (dashed lines), for the tumors in (A). Increasing the stiffness of the gel makes the tumor more compressive, and reduces the magnitude of circumferential stress outside the tumor.

where $RSS$ is the residual sum of squares, $n$ is number of observed data points, and $K$ is the number of estimable parameters in the model. The number of radius data points is 12 for the tumor in free suspension, 8 for the tumor in 0.7% agarose gel, and 8 for the tumor in 1.0% gel. So, $n = 12 + 8 + 8 = 28$.

The model without mechanical feedback has $K = 3$ parameters $(\lambda_0, \lambda_{A,0}, L)$, $RSS = 76.37^2$, and the AICc score is 156.49. The model with mechanical feedback has $K = 9$ parameters $(\lambda_0, \lambda_{A,0}, L, \beta, \Delta_A, \gamma_A, n, c_{H,0.7\%}, c_{H,1.0\%})$, $RSS = 24.19^2$, and the AICc score is 113.12. This indicates the model without feedback is $e^{(113.12-156.49)/2} = 3.8 \times 10^{-10}$ times as probable as the model with feedback.

Next, we use the previous fitting to predict the gel release data in Fig. 1B in [1]. We use the same parameters prior to gel release. The time of gel release is varied from $T = 25$ to $T = 35$, at which time we set $c_H = 0$. The time points that minimize the $l^2$-error of the model are:

$$0.7\%\text{: gel release at } T = 34$$
$$1.0\%\text{: gel release at } T = 28.$$

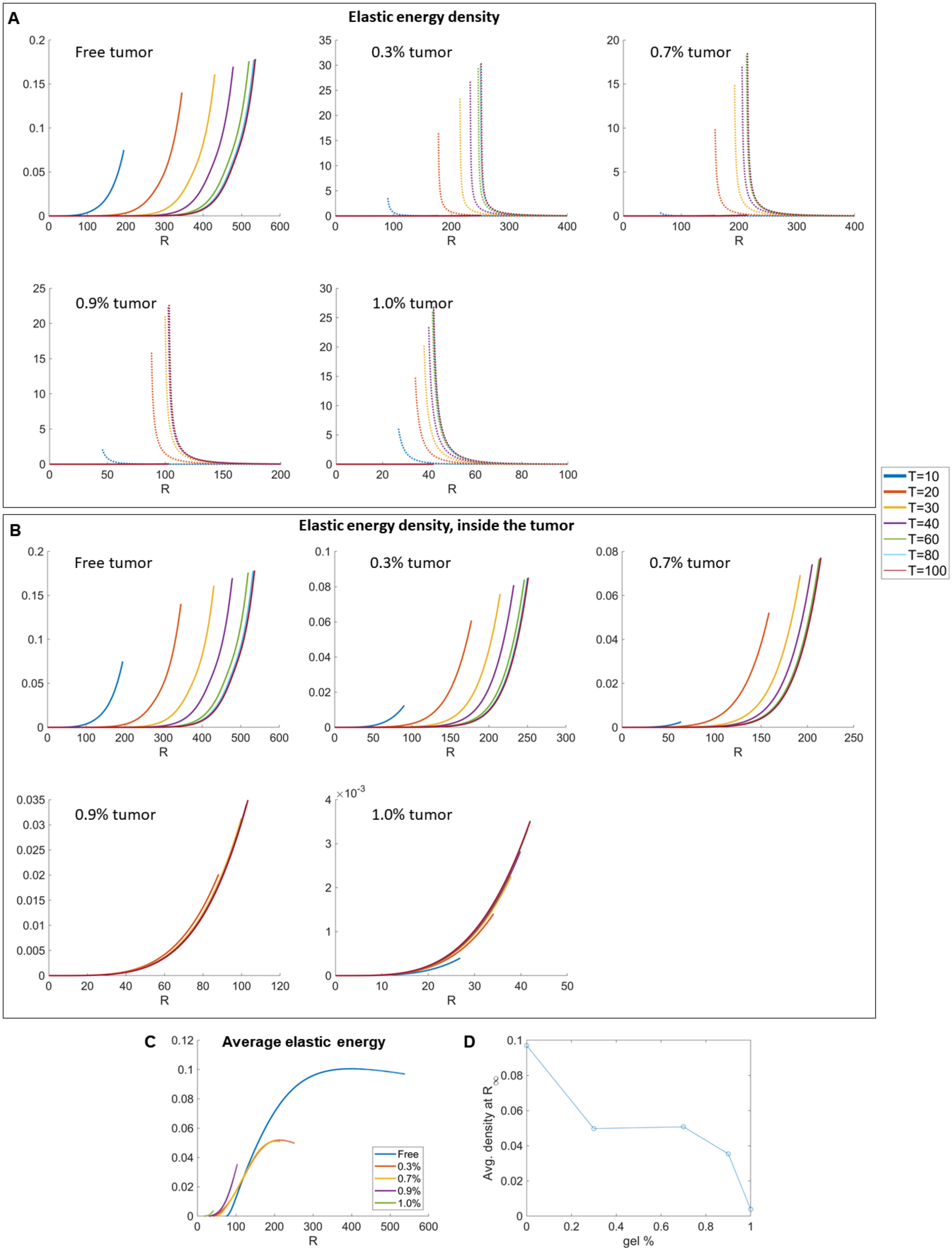


Fig.S 2: (A-F) Time evolution of the elastic energy density for the tumors in Figure 1A. The energy density is much larger outside the tumor, and is reduced by the gel stiffness. (G-L) Elastic energy density inside the tumor is reduced by the gel stiffness. (M) Spatial distribution of the average elastic density inside the tumor. (N) The elastic density at tumor boundary is reduced by the gel stiffness.

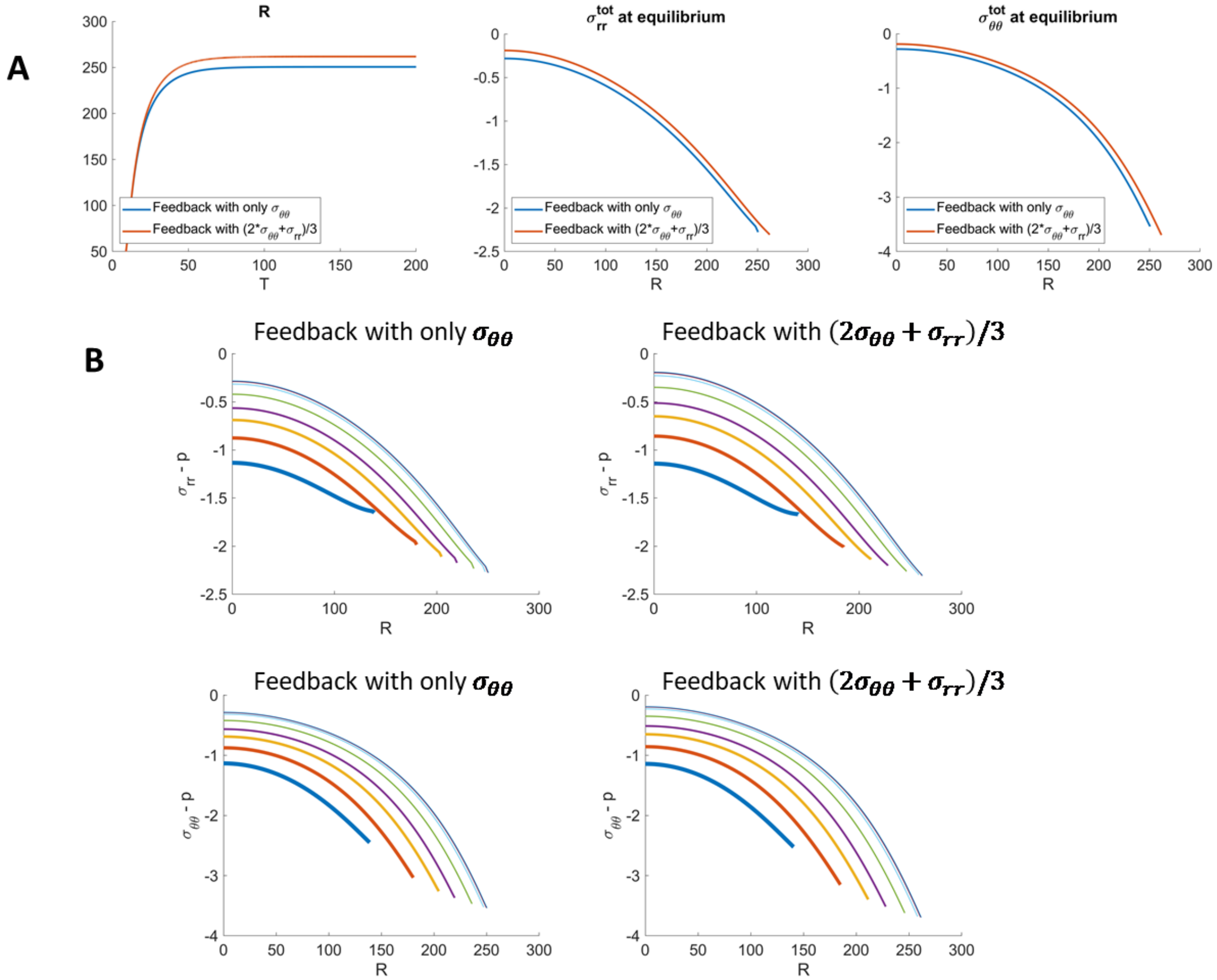


Fig.S 3: Confined tumor growth in 0.3% gel with different choices of stress-mediated feedback. (A) Evolution of tumor radius with only $\sigma_{\theta\theta}$ in the feedback (Eq. (3)), and with the stress invariant $(2\sigma_{\theta\theta} + \sigma_{rr})/3$ in the feedback (Eq. (4)). (B) Evolution of total radial stress $\sigma_{rr} - p$ and total hoop stress $\sigma_{\theta\theta} - p$ with different feedback. The approximation by $\sigma_{\theta\theta}$ does not change the results significantly.

For the above parameter fittings, we have used the feedback function Eq.(9) in the main text:

$$\lambda_E(r,t) = \Delta_A \frac{\gamma_A(\sigma_{\theta\theta}^{tot})^m \cdot 1_{\{\sigma_{\theta\theta}^{tot}<0\}}}{1+\gamma_A(\sigma_{\theta\theta}^{tot})^m \cdot 1_{\{\sigma_{\theta\theta}^{tot}<0\}}} \tag{3}$$

where only $\sigma_{\theta\theta}^{tot}$ is considered as the input. This can be considered as an approximation of the function

$$\lambda_E(r,t) = \Delta_A \frac{\gamma_A((2\sigma_{\theta\theta}^{tot}+\sigma_{rr}^{tot})/3)^m \cdot 1_{\{2\sigma_{\theta\theta}^{tot}+\sigma_{rr}^{tot})/3<0\}}}{1+\gamma_A((2\sigma_{\theta\theta}^{tot}+\sigma_{rr}^{tot})/3)^m \cdot 1_{\{2\sigma_{\theta\theta}^{tot}+\sigma_{rr}^{tot})/3<0\}}} \tag{4}$$

where $(2\sigma_{\theta\theta}^{tot} + \sigma_{rr}^{tot})/3$ is an invariant of the Cauchy stress tensor. We show in Figure S3 that the error from this approximation does not change the results significantly.

---

[1] G. Helmlinger, P. A. Netti, H. C. Lichtenbeld, R. J. Melder, and R. K. Jain, Nature biotechnology **15**, 778 (1997).
[2] K. Burnham and D. Anderson, *Model Selection and Multimodel Inference: A practical information-theoretic approach (2nd ed.)* (Springer-Verlag, 2002).
