## Supplemental Data for Figure 3 for "Stress generation, relaxation and size control in confined tumor growth"

(Dated: February 6, 2021)

### 3. SUPPLEMENTAL DATA FOR FIGURE 3

**Parameters for Figure 3**

| | |
|---|---|
| $R_0 = 75$ | tumor initial radius (in $\mu$m) |
| $L = 40$ | diffusional length of nutrient (in $\mu$m) |
| $\beta = 0.6$ | relaxation rate |
| $\lambda_0 = 1.1$ | rate of cell volume growth |
| $\lambda_{A,c} = 0.2$ | rate of apoptosis |
| $\Delta_A = 0.9$ | maximum rate of the water flux, |
| $\gamma_A = 0.1$ | sensitivity to the feedback on $\lambda_E$ |
| $\gamma_c = 0.1$ | sensitivity to the feedback on $\lambda_c$ |
| $n = 2$ | Hill coefficient of feedback on $\lambda_E$ |
| $l = 2$ | Hill coefficient of feedback on $\lambda_c$ |
| $c_H = 0$ | nondimensional shear modulus, free tumor |
| $c_H = 1.2$ | nondimensional shear modulus, 0.7% tumor |
| $c_H = 3.4$ | nondimensional shear modulus, 1.0% tumor |
