## Supplemental Data for Figure 4 for "Stress generation, relaxation and size control in confined tumor growth"

(Dated: August 30, 2021)

### 4. SUPPLEMENTAL DATA FOR FIGURE 4

We use the data from [1, 2], where colon carcinoma tumor spheroids containing mouse CT26 cell lines were grown under isotropic compression from an osmotically-induced external pressure. Similar to the previous fitting, we assume that $\bar{p} = 0$ in the free case, and only $\bar{p}$ changes in the compressed case. The parameter combination that minimizes the sum of the $l^2$-error is:

$$\lambda_0 = 1.1, L = 10, \lambda_{A,c} = 0.05, \Delta_A = 0.3, \gamma_A = 0.1, n = 2, \beta = 0.4$$
$$\bar{p} = 2.1 \text{ for the compressed tumor}$$

Without feedback from mechanical stresses, the parameter combination that minimizes the $l^2$-error is:

| | $\lambda$ | $\lambda_A$ | $L$ |
|---|---|---|---|
| Free | 1.6 | 1.2 | 80 |
| Compressed | 1.6 | 1.1 | 50 |

Recall that in Figure 2EF (in the main text), when stresses are released by removing the gels, the tumors tend to recover the growth of the unconstrained spheroid. We report similar observations in the case of osmotic pressure. When the pressure is released [1], growth inhibition is reversed and the tumor tends to regrow to the uncompressed (free) size (Figure 4EF). The parameter combination that minimizes the $l^2$-error using the data presented in Fig. 1 in [1] is:

$$\lambda_0 = 2.1, L = 130, \lambda_{A,0} = 1.4, \Delta_A = 0.4, \gamma_A = 0.2, n = 2, \beta = 1.8$$
$$\bar{p} = 0.3 \text{ for the compressed tumor}$$
$$\text{pressure release at } T = 12$$

We list the fitted parameter values of the data in [1–3] below:

| | Description | Fig. 2,3 [3] | Fig. 4A-4D [1] | Fig. 4E-4F [2] |
|---|---|---|---|---|
| $\lambda_0$ | rate of cell volume growth (per day) | 1.1 | 1.1 | 2.1 |
| $L$ | diffusional length of nutrient ($\mu$m) | 40 | 10 | 130 |
| $\lambda_{A,c}$ | rate of apoptosis (per day) | 0.2 | 0.05 | 1.4 |
| $\Delta_A$ | maximum rate of the water flux (per day) | 0.9 | 0.3 | 0.4 |
| $\gamma_A$ | sensitivity to the feedback on $\lambda_E$ | 0.1 | 0.1 | 0.2 |
| $n$ | Hill coefficient of feedback on $\lambda_E$ | 2 | 2 | 2 |
| $\beta$ | relaxation rate (per day) | 0.6 | 0.4 | 1.8 |
| $c_H$ | nondimensional shear modulus, free tumor | 0 | | |
| $c_{H,0.7\%}$ | nondimensional shear modulus, 0.7% tumor | 1.2 | | |
| $c_{H,1.0\%}$ | nondimensional shear modulus, 1.0% tumor | 3.4 | | |
| $\bar{p}_f$ | nondimensional external pressure, free tumor | | 0 | 0 |
| $\bar{p}_c$ | nondimensional external pressure, compressed tumor | | 2.1 | 0.3 |

Note that the data in [2] (Figs. 4E-4F) is different from that in [1] (Figs. 4A-D) and correspondingly the fitted parameters are different. In particular, the tumor spheroids from [2] grow more rapidly at early times than those from [1]. Therefore, the rate of cell volume growth ($\lambda_0$), rate of apoptosis ($\lambda_{A,c}$), and diffusional length of nutrient ($L$) are much larger for Figs. 4E-F than those for Figs. 4A-D.

In Figure 3 in the main text, we showed that upon gel removal, the spatial distribution of the cell density ($\rho_c$) reverse to the distribution in the free-boundary case. In the case of isotropic compression from an osmotically-induced external pressure ([1]), the distribution of $\rho_c$ also reverses to that in the free tumor.

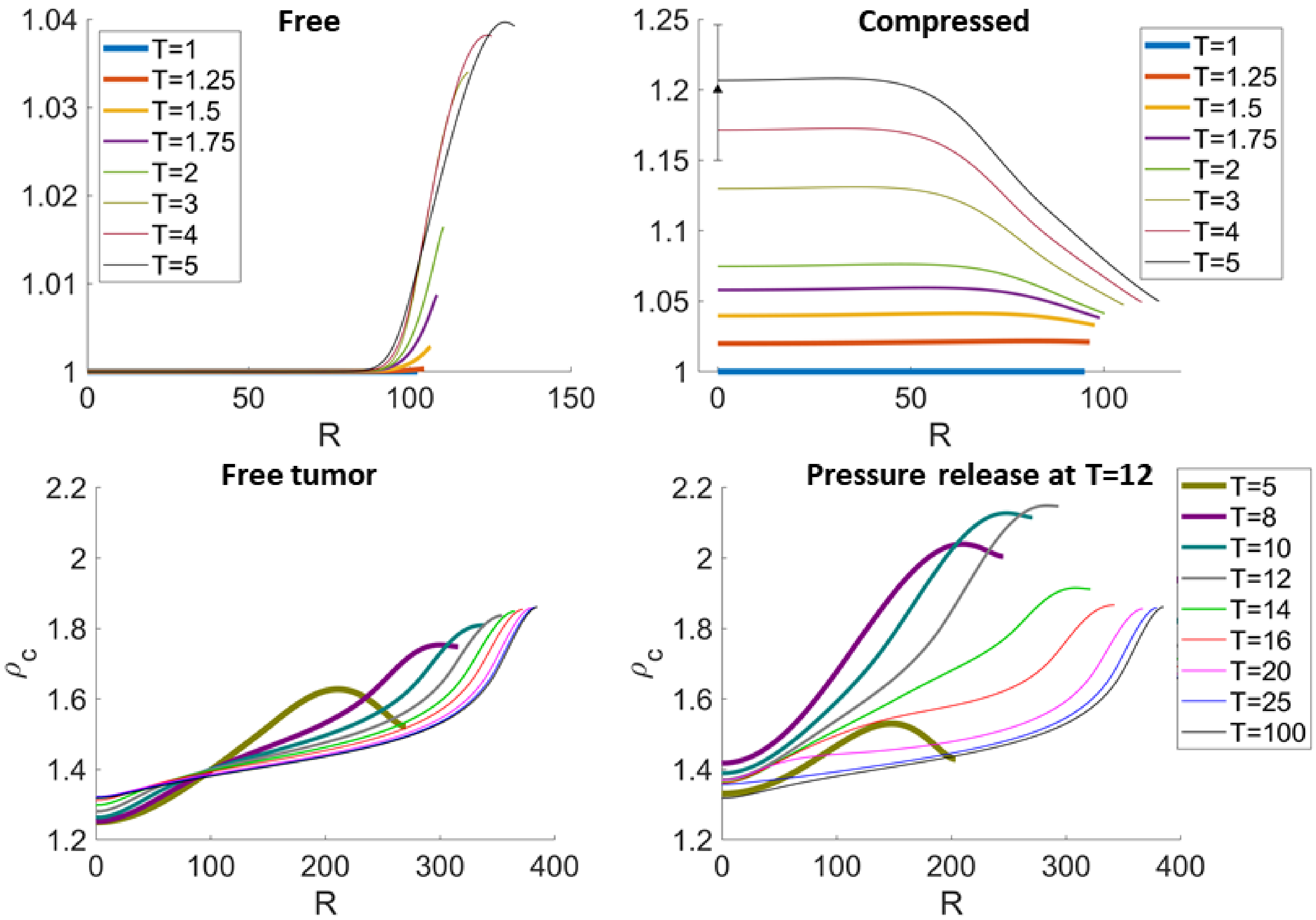


Fig.S 1: Top row: cell densities $\rho_c$ in the free and constrained tumor in Figure 4A. Data in [2] suggests that the cell density at tumor center is approximately 20% larger in the compressed tumor, compared to the free tumor. This is shown by the error bar in the right panel. Similar with Figure 3, the regions with largest cell density shifts towards the tumor center, indicating cells are more packed inside the tumor. Bottom row: after releasing the external pressure, the cell density of the compressed tumor (right panel) recovers to the same level in the free tumor (left panel).


[1] F. Montel, M. Delarue, J. Elgeti, L. Malaquin, M. Basan, T. Risler, B. Cabane, D. Vignjevic, J. Prost, G. Cappello, et al., Physical review letters **107**, 188102 (2011).

[2] M. Delarue, F. Montel, D. Vignjevic, J. Prost, J.-F. Joanny, and G. Cappello, Biophysical journal **107**, 1821 (2014).

[3] G. Helmlinger, P. A. Netti, H. C. Lichtenbeld, R. J. Melder, and R. K. Jain, Nature biotechnology **15**, 778 (1997).
