## Supplemental Data for Figure 5 for "Stress generation, relaxation and size control in confined tumor growth"

(Dated: February 18, 2021)

### 5. SUPPLEMENTAL DATA FOR FIGURE 5

The effect of $\beta$ on tumor radius is shown in Figure S1. $R_\infty$ is increased by $\beta$, which reduces the feedback by reducing the compressive stress at the tumor boundary (see Figure S5 in S1 Text). When $\beta$ is small, the tumor radius overshoots before reaching steady state (e.g. blue curve in Figure S1), because the circumferential stress reaches steady state faster than the radius. Details of the radii, stresses, elastic energy, and energy density distributions are shown in Figure S2-S6.

**Parameters for Figure 5**

| | |
|---|---|
| $R_0 = 75$ | tumor initial radius (in $\mu$m) |
| $L = 40$ | diffusional length of nutrient (in $\mu$m) |
| $\lambda_0 = 1.1$ | rate of cell volume growth |
| $\lambda_{A,c} = 0.2$ | rate of apoptosis |
| $\Delta_A = 0.9$ | maximum rate of the water flux, |
| $\gamma_A = 0.1$ | sensitivity to the feedback on $\lambda_E$ |
| $\gamma_c = 0.1$ | sensitivity to the feedback on $\lambda_c$ |
| $n = 2$ | Hill coefficient of feedback on $\lambda_E$ |
| $l = 2$ | Hill coefficient of feedback on $\lambda_c$ |

$\beta = 0.05, 0.1, 0.2, 0.4, 0.6, 0.8, 1$ relaxation rate

$c_H = 0, 0.1, 0.2, 0.3, \cdots, 1$ nondimensional shear modulus

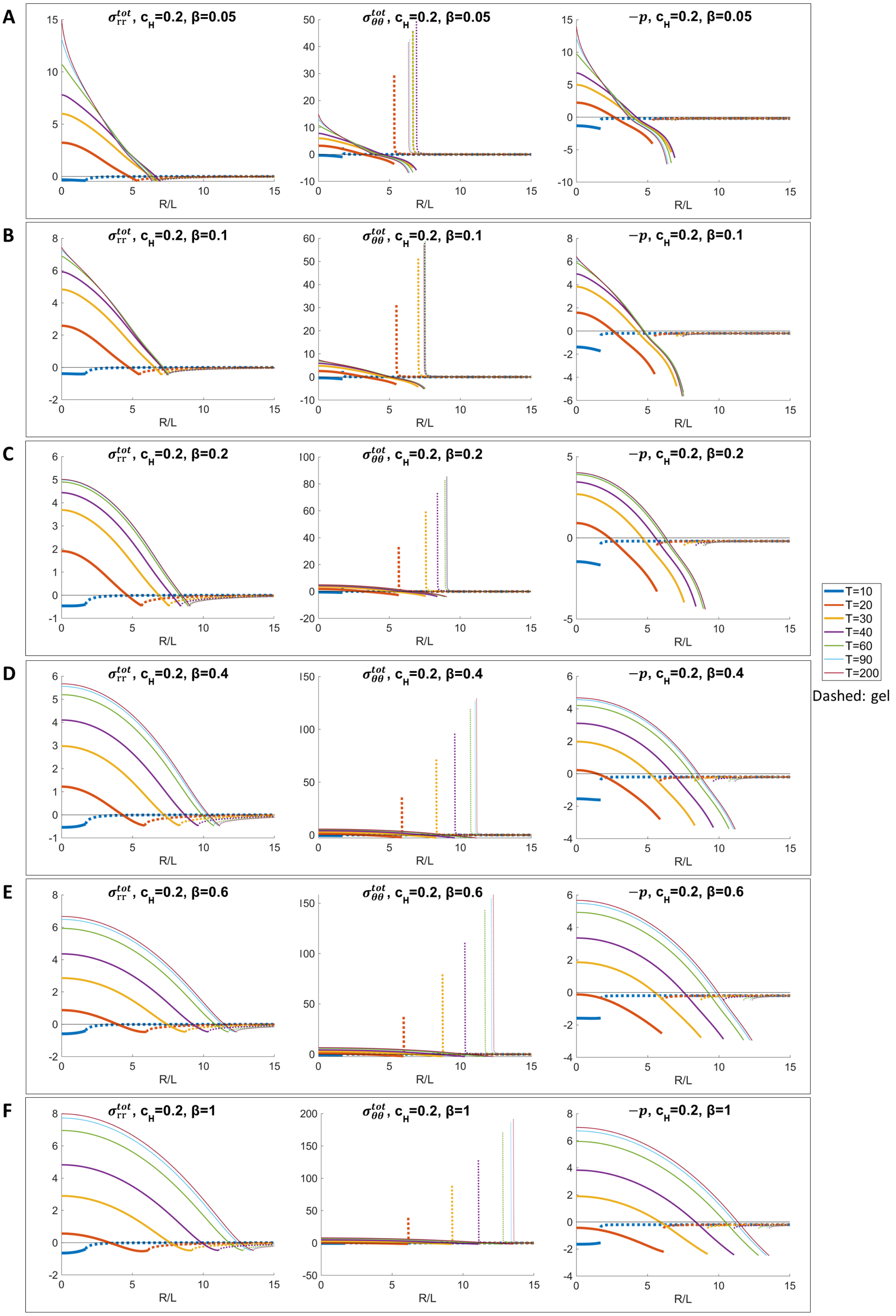


Fig.S 1: Time evolution of the total radial, circumferential stress and pressure inside the tumor (solid lines) and outside (dashed lines), with $c_H = 0.2$ and different $\beta$. As $\beta$ increases from 0.05 to 0.2 (A-C), the magnitude of tensile stress in the tumor is reduced by stress relaxation. When $\beta \geq 0.4$ (D-F), the magnitude of tensile stress increases with $\beta$ as a size effect of larger tumor.

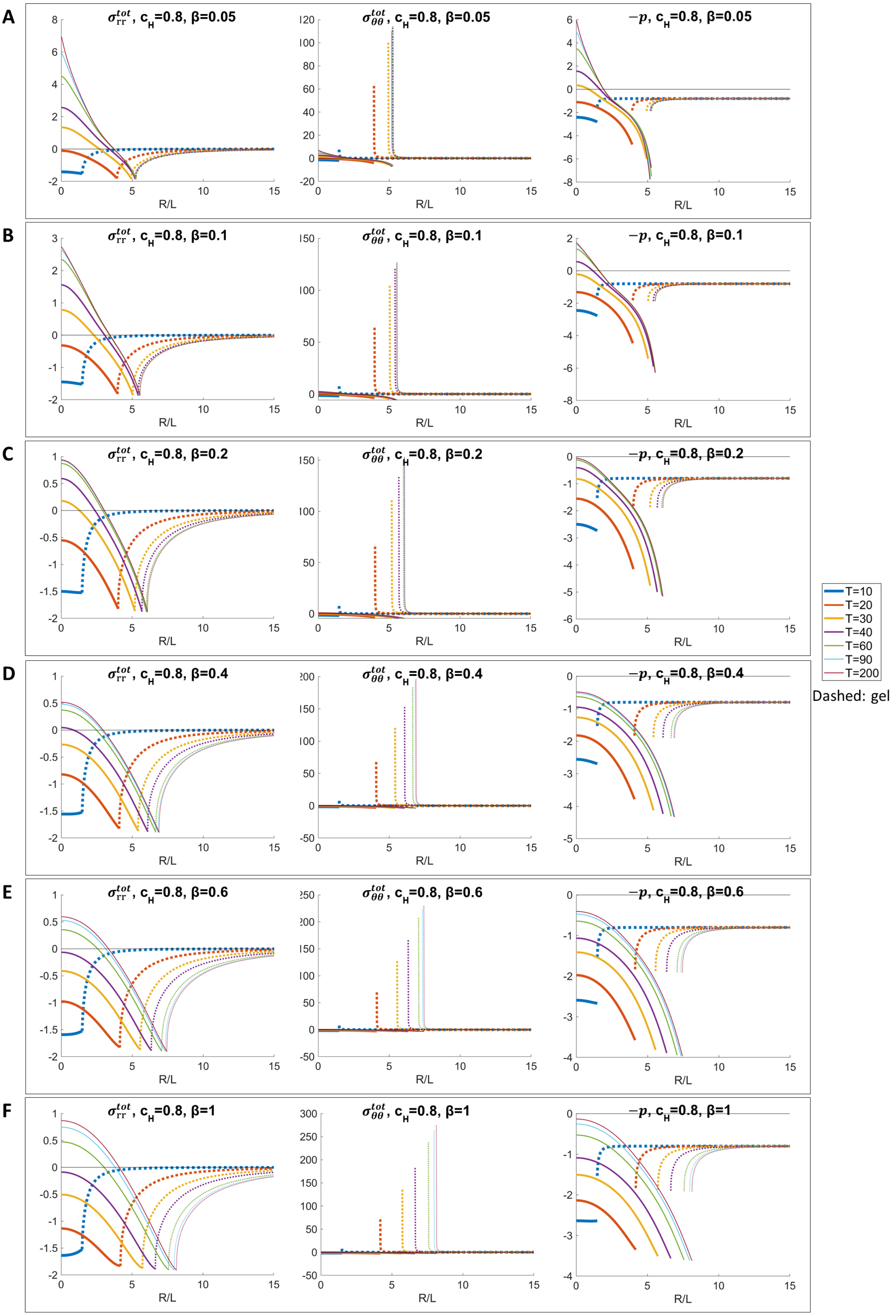


Fig.S 2: Time evolution of the total radial, circumferential stress and pressure inside the tumor (solid lines) and outside (dashed lines), with $c_H = 0.8$ and different $\beta$. The tumor is tensile inside when $\beta = 0.05$. The tumor becomes more compressive as $\beta$ increases. The stress outside the tumor is determined by $c_H$ and $R_\infty$ (Eq. 63 in S1 Text).

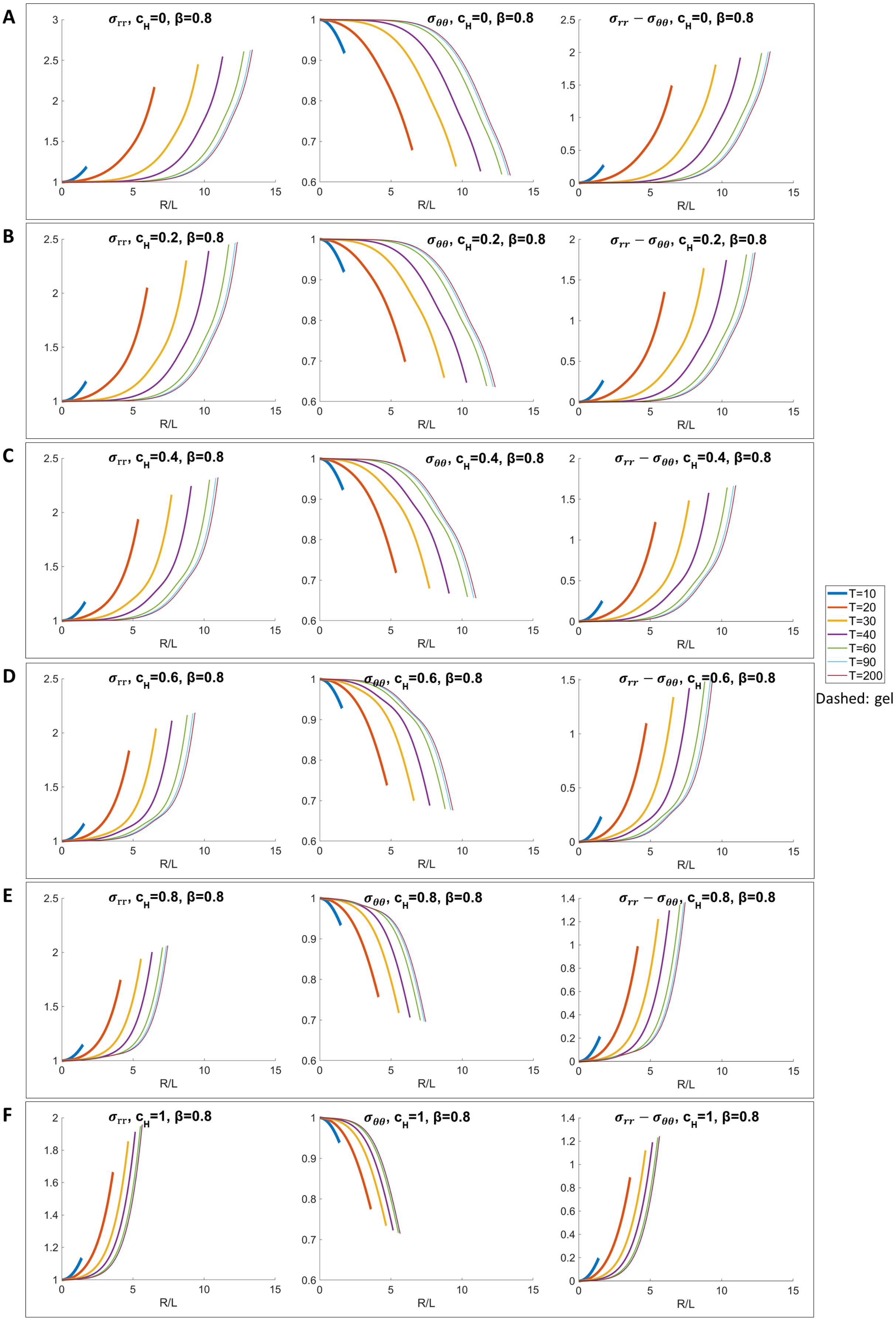


Fig.S 3: Plots of the radial stress ($\sigma_{rr}$), circumferential stress ($\sigma_{\theta\theta}$), and stress anisotropy ($\sigma_{rr} - \sigma_{\theta\theta}$) with different $c_H$. Larger $c_H$ reduces the magnitude and results in more homogeneous stress.

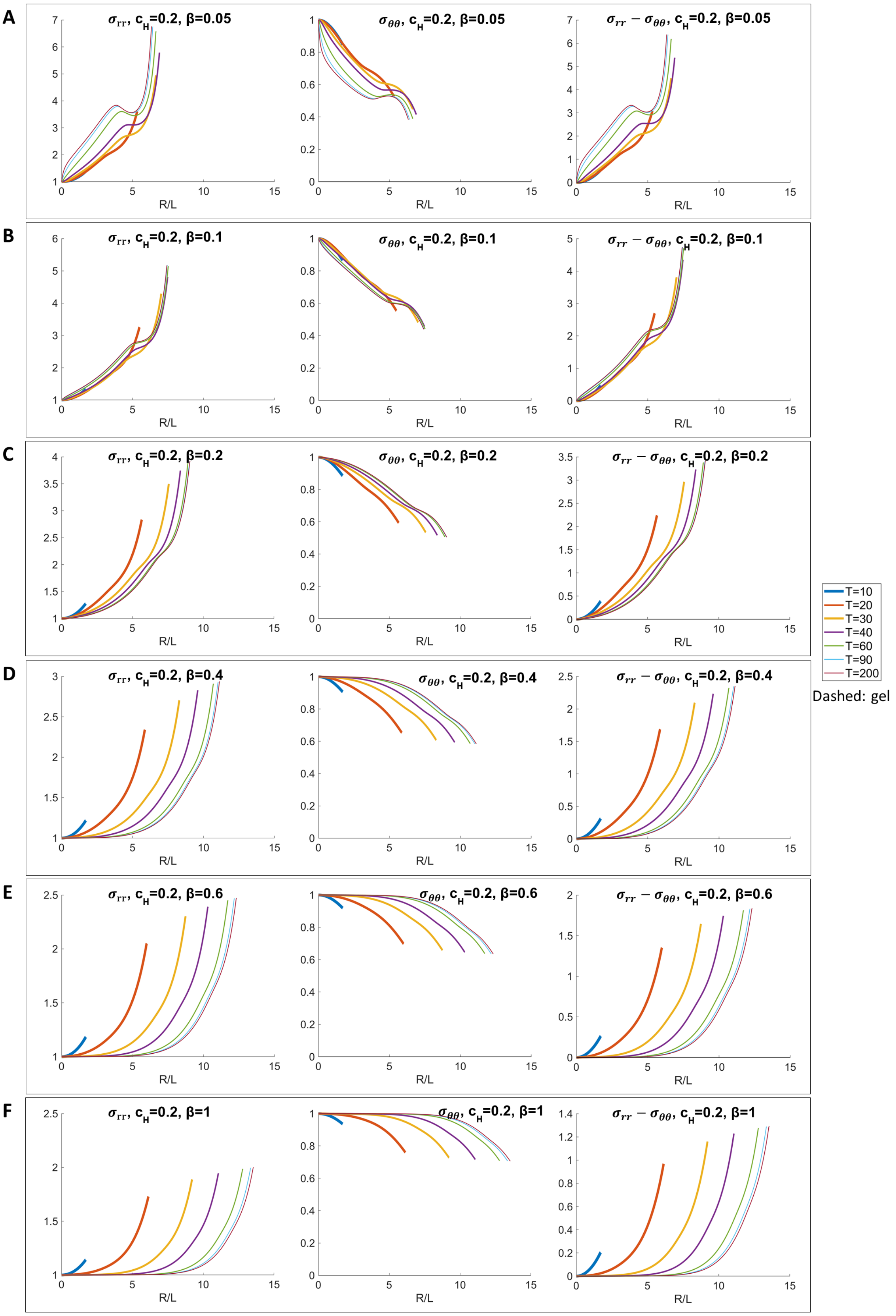


Fig.S 4: Plots of the radial stress ($\sigma_{rr}$), circumferential stress ($\sigma_{\theta\theta}$), and stress anisotropy ($\sigma_{rr} - \sigma_{\theta\theta}$) with $c_H = 0.2$ and different $\beta$. Larger $\beta$ changes the shape of stress, reduces the magnitude and results in more homogeneous stress.

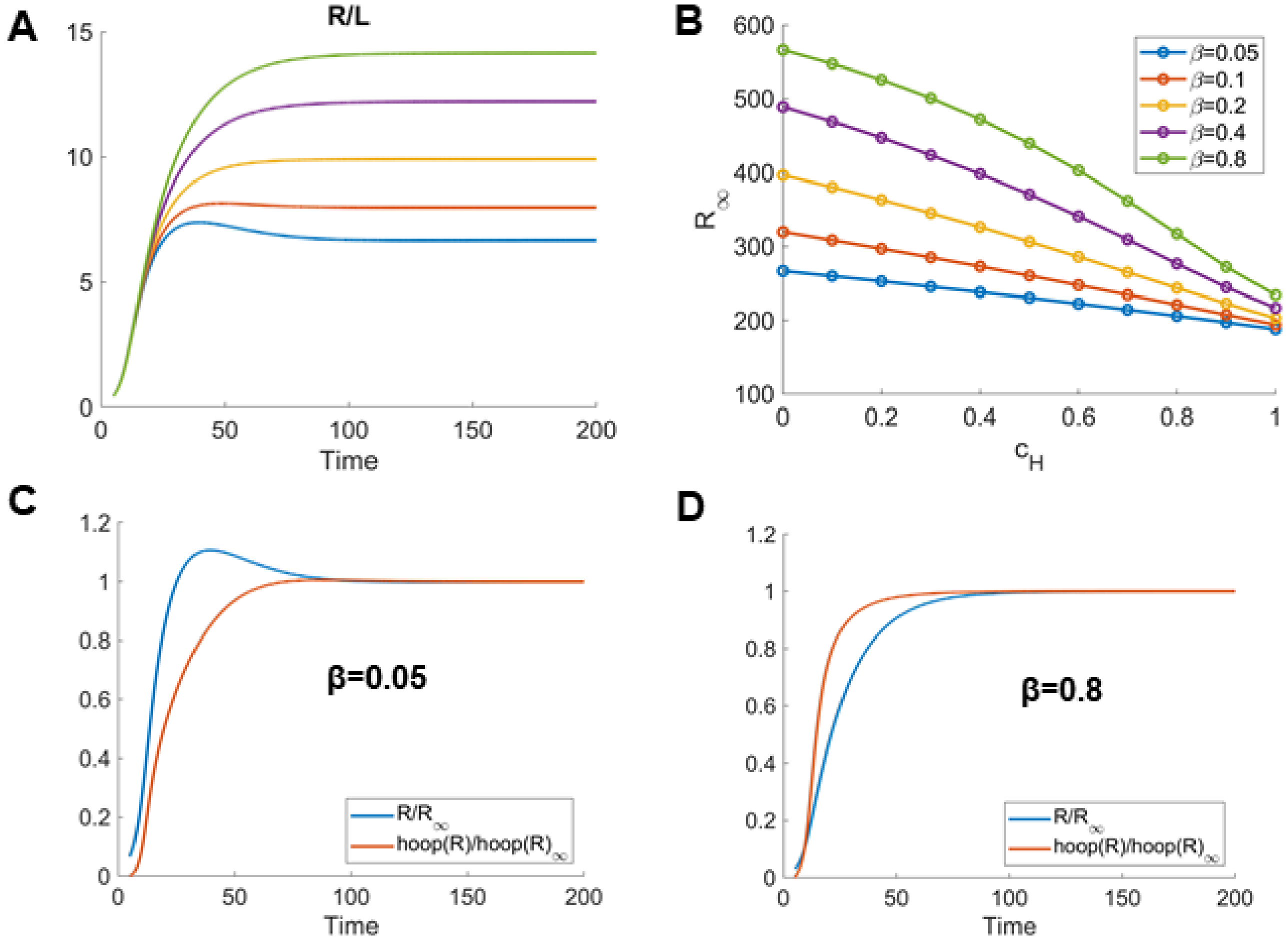


Fig.S 5: (A) Evolution of the tumor radius with $c_H = 0$ and different $\beta$. The steady state size is increased by $\beta$, which reduces the compressive stress at the tumor boundary. Consequently, the feedback on the apoptosis rate is smaller. When $\beta$ is small (e.g. 0.05), the tumor radius overshoots before reaching steady state, corresponding to the parameter region marked by the red dashed curves in Figure 5A in the main text. (B) Tumor radius at equilibrium as a function of $c_H$, for different $\beta$. (C, D) The spheroid radius and circumferential (hoop) stress, relative to their values at equilibrium, as a function of time for $\beta = 0.05$ (C) and $\beta = 0.8$ (D), with $c_H = 0$.
